# The performance of deep generative models for learning joint embeddings of single-cell multi-omics data

**DOI:** 10.1101/2022.06.06.494951

**Authors:** Eva Brombacher, Maren Hackenberg, Clemens Kreutz, Harald Binder, Martin Treppner

## Abstract

Recent extensions of single-cell studies to multiple data modalities raise new questions regarding experimental design. For example, the challenge of sparsity in single-omics data might be partly resolved by compensating for missing information across modalities. In particular, deep learning approaches, such as deep generative models (DGMs), can potentially uncover complex patterns via a joint embedding. Yet, this also raises the question of sample size requirements for identifying such patterns from single-cell multi-omics data. Here, we empirically examine the quality of DGM-based integrations for varying sample sizes. We first review the existing literature and give a short overview of deep learning methods for multi-omics integration. Next, we consider eight popular tools in more detail and examine their robustness to different cell numbers, covering two of the most common multi-omics types currently favored. Specifically, we use data featuring simultaneous gene expression measurements at the RNA level and protein abundance measurements for cell surface proteins (CITE-seq), as well as data where chromatin accessibility and RNA expression are measured in thousands of cells (10x Multiome). We examine the ability of the methods to learn joint embeddings based on biological and technical metrics. Finally, we provide recommendations for the design of multi-omics experiments and discuss potential future developments.

## 1 Introduction

Many diseases, such as cancer, affect complex molecular pathways across different biological layers. Consequently, there is currently an ongoing surge in multi-omics techniques that study the interaction of biomolecules across various omics layers [47, 59]. Multi-omics techniques have been used, e.g., to infer mechanistic insights about molecular regulation, the discovery of new cell types, and the delineation of cellular differentiation trajectories [1, 13, 54, 58]. However, because performing multiomics experiments in the same cell is still costly and experimentally complex, many experiments have been carried out with comparatively small numbers of cells so far. Additionally, single-cell multi-omics data suffer from the sparseness and noisiness of the measured modalities, differences in sequencing depth, and batch effects. Data analysis is further complicated by differing feature spaces as well as shared and modality- or batch-specific variation [33].

Deep learning approaches, known for their ability to learn complex non-linear patterns from data, have become a popular building block for integrating different data types [17, 22]. For example, in 2021’s Conference on Neural Information Processing Systems (NeurIPS) competition (https://openproblems.bio/neurips_2021), which addressed the topic of multimodal single-cell data integration, neural networks proved to be the most popular model choice, with shallow deep learning models being among the best-performing methods [33]. Specifically, deep generative models (DGMs), such as variational autoencoders (VAEs), are increasingly employed to infer joint embeddings, i.e., low-dimensional representations, from multi-omics datasets. This allows for performing all further downstream analyses simultaneously within this joint latent space (**Figure 1 B**).

**Figure 1:**
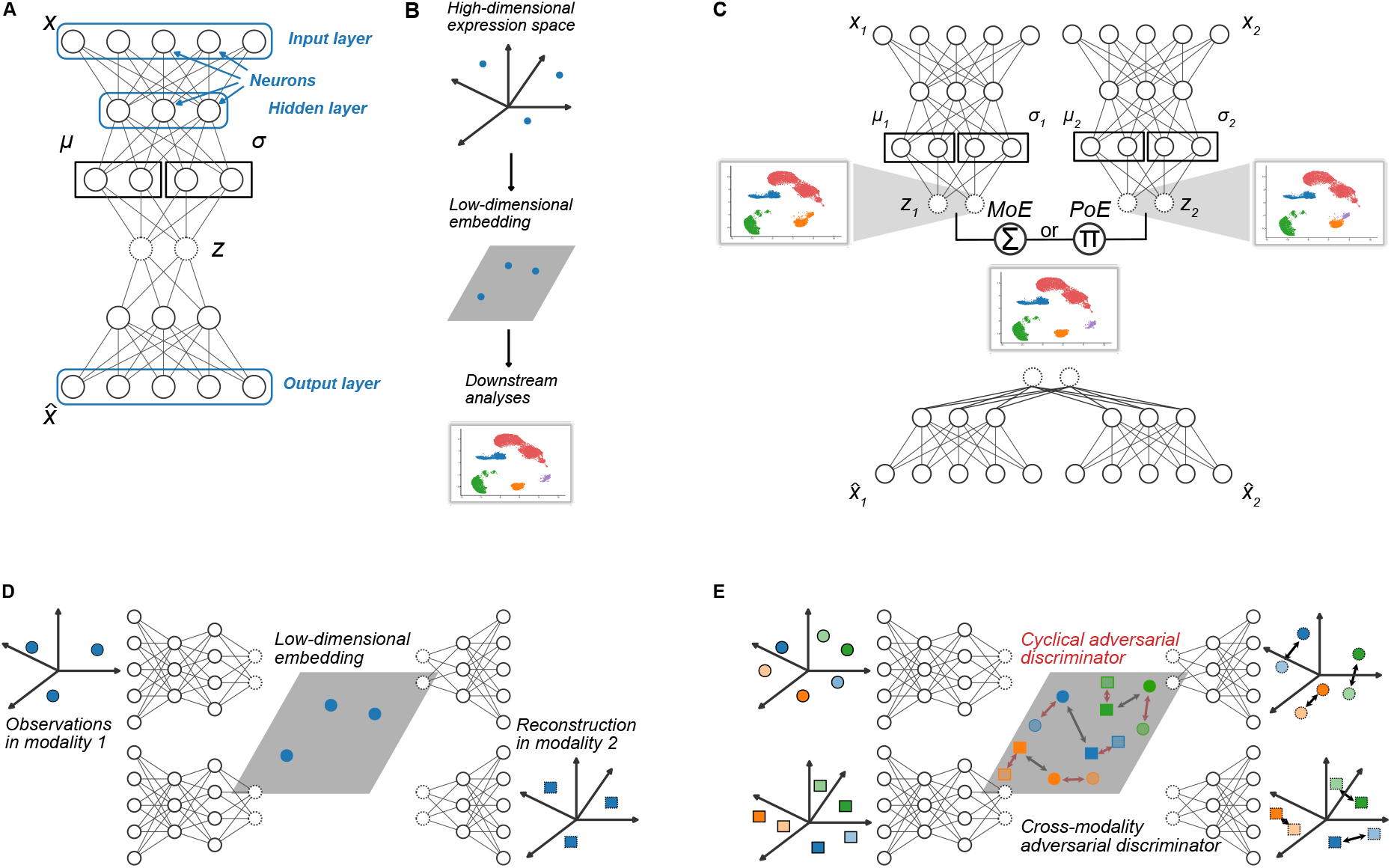
Neural network architectures. (A) Exemplary network architecture of vanilla VAE, where *x* represents the input data and 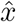 is the reconstructed data. Random variables *z* in the bottleneck layer are indicated by dashed circles. *μ* and *σ* represent the mean and standard deviation of the distributions, typically Gaussian distributions with diagonal covariance matrices, learned in the bottleneck layer. (B) Typical workflow: High-dimensional omics data are mapped to a low-dimensional embedding, which can then be utilized for visualization and downstream analyses such as clustering or trajectory inference. (C) General architecture of multimodal VAEs. (D) Cross-modality translation: High-dimensional measurements from one modality are mapped to a low-dimensional embedding with the modalityspecific encoder. The latent representation is then used as input for the decoder of the respective other modality. (E) Adversarial training principles: Adversarial discriminators can be employed (1) to align low-dimensional embeddings of different modalities (squares vs. circles) of the same cell (same color) in the latent space (black arrows), (2) to align reconstructed profiles with the cross-modal reconstructions (lighter colors) obtained by decoding low-dimensional embeddings of one modality with the decoder of the other modality (black arrows in the reconstruction space, or (3) to align re-embedded decoder outputs from intra-modal and/or cross-modal reconstruction (lighter colors) with the original embeddings (red arrows in the latent space).

This review provides a systematic overview of current DGM-based approaches for learning joint embeddings from multi-omics data and illustrates how small sample sizes impact the amount of information that can be recovered from multi-omics datasets. Specifically, we examine how the performance of popular DGM-based approaches to infer joint low-dimensional representations from such data is influenced by varying numbers of cells. The required number of cells is particularly relevant at the stage of designing an experiment [56]. To tackle the challenging task of evaluating the quality of a latent representation with respect to the conservation of biological signal and batch correction capabilities, we draw on the guidelines provided by Luecken et al. [40].

The training of DGMs on multi-omics data is challenging due to the inherent high dimensionality and low sample size of multi-omics data and the large number of model parameters that need to be estimated while avoiding overfitting and bias [25]. Thus, we investigate the impact of cell numbers on the performance of selected single-cell multi-omics integration algorithms. We consider eight popular VAE-based tools that incorporate different integration paradigms and training strategies for this illustration. Specifically, we included product-of-experts- and mixture-of-experts-based approaches and techniques that employ additional, commonly used integration techniques, such as cross-modality translation and adversarial training. Also, we chose models with different degrees of architectural complexity, including one model [34] with (self-)attention modules and additional regularization by clustering consistency. We thus created an exemplary selection that represents the range of architectural choices, additional training and regularization strategies, and levels of complexity currently used for the task at hand. Thus, viewing the selected models as representatives of the current landscape of DGMs for multi-omics integration, our case study enables us to draw conclusions on the performance of the investigated tools in small sample size scenarios, and to give recommendations regarding architectural choices, integration strategies, and regularization paradigms.

## 2 Deep learning background

As the number of experimental methods in molecular biology is exploding, immense amounts of data are produced. Machine learning techniques can help in extracting information from such data to make it human-interpretable.

In recent years, deep learning has emerged as a potent tool for analyzing such high-throughput biological data. At the core of these approaches are artificial neural networks (ANNs) that provide powerful yet versatile building blocks to learn complex non-linear transformations and thus uncover underlying structures from high-dimensional data.

In particular, a networks’ architecture comprises interconnected layers of neurons. Each neuron is connected to all of the neurons in the preceding layer. The depth of the network is determined by the number of hidden layers, i.e., the layers between the input and output layers. In contrast, the number of neurons in one layer determines a network’s width (**Figure 1 A**). With deep architectures, ANNs are especially effective at learning increasingly complicated patterns from large volumes of data based on non-linear transformations. Specifically, each individual neuron computes a weighted sum of its inputs, where the weighted total is then subjected to an activation function, typically producing a nonlinear transformation of the neuron’s output. The weights of an ANN, which link the neurons between layers and make up the model’s parameters, are a crucial part of the model. Training an ANN amounts to finding model weights that optimize a loss function, which represents how well the model fits the data. However, one of the major difficulties in training ANNs is optimizing the loss function as it is typically complex and non-convex and the parameter space is high-dimensional [4].

While supervised deep learning relies on labeled data to solve, e.g., classification problems, unsupervised deep learning can be employed in exploratory analyses to uncover central structure in data. For example, researchers frequently aim to understand cell-type compositions, for which they usually rely on unlabelled data. Hence, unsupervised deep learning methods have become increasingly popular in omics data analysis. Specifically, DGMs have been used for imputation [38, 67], visualization of the underlying structure of single-cell RNA-sequencing (scRNA-seq) data [16], and synthetic data generation [44, 56].

Many computational approaches for processing scRNA-seq data use dimensionality reduction to produce a compressed representation of the high-dimensional transcription space. Grouping cells based on some measure of distance is a typical step in scRNA-seq research since these analyses usually attempt to understand the cell type composition of tissues or samples. However, conventional distance metrics, such as Euclidean distance, are unsuited to accurately represent similarity relations between cells due to the high dimensionality of the gene expression space, which is commonly referred to as the curse of dimensionality. As a result, the solution usually adopted is to reduce the number of dimensions based on the assumption that such a low-dimensional space captures the underlying biological phenomena. As an illustration, a transcription factor may be responsible for the activation of many genes. Therefore, one variable characterizing the activation of genes through the transcription factor would be adequate to describe the patterns of gene expression rather than modeling the high-dimensional space spanned by all genes and their combinations [27]. Principal component analysis (PCA) is one method for reducing the dimensionality of scRNA-seq data. However, applying PCA to scRNA-seq data has a number of drawbacks since it assumes a symmetric distribution, which is typically not satisfied in scRNA-seq data, and only learns linear relationships. As a result, researchers have developed DGMs that accurately represent the distributional assumptions of scRNA-seq data while accurately portraying the data’s inherent complexity [23, 38].

An autoencoder is the basis for many DGMs and is composed of three modules: an encoder, a bottleneck layer, and a decoder. The encoder reduces the input to a lower dimension (through the bottleneck layer), and the decoder reconstructs the original input from the bottleneck. This design also forms the foundation for the variational autoencoder and effectively compresses the essential information needed for data reconstruction [37], which is mainly used to eliminate noise from data by compressing and re-compressing and reducing data to lower dimensions for visualization. In contrast, a variational autoencoder aims to infer the parameters of the probability distribution assumed to underly the source data, which can subsequently be used to generate realistic in-silico data.

Specifically, DGMs are trained to capture the joint probability distribution over all features in the input data, thus allowing to also generate new synthetic data with the same patterns as the training data by sampling from the learned distribution. This is typically done by introducing latent random variables *z* in addition to the observed data *x*. In single-cell transcriptomics applications, these latent variables might encode complex gene programs based on non-linear relationships between genes. Typically, the joint distribution *p_θ_* (*x, z*) of observed and latent variables is described through a parametric model, where *θ* represents the model parameters. The joint probability can be factorized into a prior probability *p_θ_*(*z*) and a posterior *p_θ_*(*x*|*z*) and can thus be written as *p_θ_* (*x,z*) = *p_θ_*(*z*)*p_θ_*(*x*|*z*). Inferring the data likelihood *p_θ_*(*x*) = ∫ *p_θ_*(*x, z*)*dz* from the joint distribution requires marginalizing over all possible values of z, which is typically computationally intractable [30]. Hence, approximate inference techniques are employed to efficiently optimize the model parameters [9].

Two methods are frequently used in the machine learning literature to aggregate distributions, such as data from various single-cell modalities like gene expression and surface proteins. One strategy involves multiplying the density functions of the two modalities to create a product of experts (PoE) approach. On the other side, a mixture of experts (MoE) approach can blend the modalities using a weighted sum. In Section 2.2, we go over these strategies’ benefits and drawbacks.

In single-cell applications, the most frequently used DGMs to date are Variational autoencoders (VAEs) [29] and generative adversarial networks (GANs) [21], which we present in more detail below.

### 2.1 Variational autoencoders

VAEs employ two independently parameterized but jointly optimized neural network models to learn an explicit parametrization of the underlying probability distributions. This is achieved by non-linearly encoding the data into a lower-dimensional latent space and reconstructing back to the data space. Specifically, the encoder (or recognition model) maps the input data x to a lower-dimensional representation given by a sample of the latent variable z, while the decoder network performs a reverse transformation and aims to reconstruct the input data based on the lower-dimensional latent representation (**Figure 1 A**).

To approximate the underlying data distribution *p_θ_* (x), the encoder and decoder parameterize the conditional distributions *p_θ1_*(*z*|*x*) and *p_θ_*(*x*|*z*), respectively. Since *p_θ_*(*x*) and *p_θ_*(*z*|*x*) are intractable, a variational approximation *q_φ_*(*z*|*x*) is employed, typically given by a Gaussian distribution with diagonal covariance matrix.

Intuitively, the model is trained by reconstructing its inputs based on the lower-dimensional data representation, such that the latent space recovers the central factors of variation that allow for approximating the data distribution as closely as possible. Formally, a training objective for the model can be derived based on variational inference [9]. The parameters *ϕ* and *θ* of the encoder and decoder distributions can be optimized by maximizing the evidence lower bound (ELBO), a lower bound for the true data likelihood *p_θ_* (*x*), with respect to *ϕ* and *θ*. Denoting with KL the Kullback-Leibler divergence KL 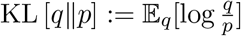 for probability distributions *q* and *p*, the ELBO is given by

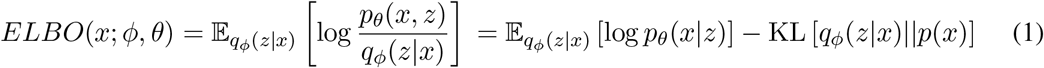

Here, the likelihood of a single observation (i.e., cell) *x* indicates how well it is supported by the model. The first term on the right side of Equation 1 describes the reconstruction error indicating how well the generated samples from the model resemble the input. The KL-divergence on the right-hand side quantifies the difference between the approximate posterior to the true posterior, and, therefore, defines the tightness of the bound—meaning the difference between the ELBO and the marginal likelihood.

The decoder network is typically built to learn the parameters of specific distributions, which best describe the underlying biological data. For scRNA-seq and surface protein data (CITE-seq) a negative binomial distribution is frequently assumed, while single-cell assay for transposase-accessible chromatin using sequencing (scATAC-seq) data usually requires an additional modeling term that accounts for the increased sparsity of the data, e.g., in the form of a zero-inflated negative binomial (ZINB) distribution [45]. Other approaches use a binarized version of the scATAC-seq data [5, 62, 68, 73].

The typical workflow for analyzing high-dimensional (single- or multi-) omics data with a VAE is illustrated in Figure 1 B. The data is embedded with the encoder to obtain a low-dimension representation, which can subsequently be used for downstream analysis, such as clustering or trajectory inference.

### 2.2 Multimodal variational autoencoders

Several approaches already exist in which multimodal VAEs [50] are used to map different omics measurements into a common latent representation [20, 39, 45]. Each of these methods uses different approaches to combine the latent variables of the respective modalities. We can usually distinguish between MoE and PoE models (**Figure 1 C**). Hence, we describe a MoE and a PoE model in more detail below and examine their performance in our analyses.

We denote a single-cell multimodal dataset as *x*_1:*M*_, where two modalities (*M* = 2) is the most common case. The joint generative model can therefore be written as 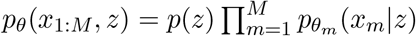, where *p_θ_m__* (*x_m_*|*z*) represents the likelihood of the decoder network for modality *m*, and *θ* = {*θ*_1_,…, *θ_M_* }.

For the MoE model, the resulting joint variational posterior can be factorized into 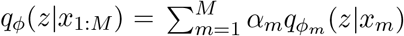, with *α_m_* = 1/*M* and *ϕ* = {*ϕ*_1_,…,*ϕ_M_*}. This results in the following ELBO:

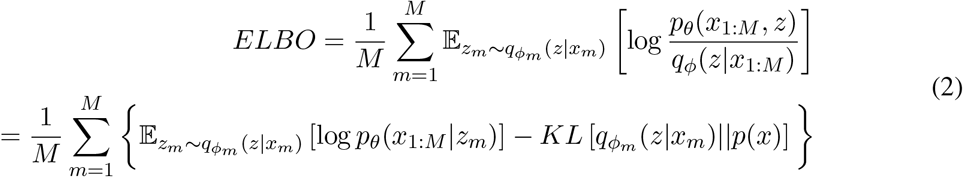

which is similar to Equation 1, but the ELBOs of the individual modalities are combined by a weighted average. In contrast, PoE approaches [20, 39] combine the variational posteriors of the individual modalities as products 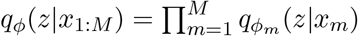.

Shi et al. [50] argue that PoE approaches suffer from potentially overconfident experts, i.e., experts with lower standard deviations will tend to have a more considerable influence on the combined posterior, as experts with lower precision come with lower marginal posteriors. In contrast, in the MoE approach we consider here, both modalities receive equal weighting, reflecting the assumption that both modalities are of similar importance. Intuitively, employing a PoE approach corresponds to taking the ‘intersection’ of the individual posteriors, as a single posterior assigning a near-zero likelihood to a specific observation is enough to cause the product to be near-zero. In contrast, an MoE approach corresponds to taking the ‘union’ of all posteriors. Additionally, the weights *α_m_* assigned to each modality can be adjusted to reflect prior assumptions on their relative importance or be learned from the data during training.

### 2.3 Cross-modality translation

In addition to architectural choices regarding the integration of the modality-specific sub-networks via a PoE or MoE approach, many VAE-based methods introduce training objectives that facilitate specific functionality such as cross-modality translation or encourage particular properties of the embedding, such as clustering consistency between the modality-specific latent representations. On a higher level, these components can be seen as regularizers that push the embeddings found by the model towards certain desired properties.

A prominent example of such an additional feature to direct a joint embedding is cross-modality translation. Here, a cell’s measurements of one modality, say, gene expression, are mapped to the joint latent space with the respective modality-specific encoder. Then, the decoder of another modality, say, chromatin accessibility, is employed to map the latent representation of the gene expression profile to a corresponding chromatin accessibility profile (Figure 1 D). This is only possible due to the integration of both modalities into a shared latent space, in which one cell’s encoded representations of different modalities align.

When paired measurements of both modalities in the same cell are available, the translated reconstructions in the respective other modality can be compared to the cell’s observed profile during training. The model learns a latent embedding that facilitates consistent cross-modality predictions. Thus, the model is explicitly pushed towards an embedding from which both modality-specific profiles can be reconstructed equally well, and that can, therefore, help in better capturing general underlying biological cell states as defined by the interplay of both modalities.

After training, cross-modality translation can be used to impute measurements of cells for which a specific modality is missing or to answer counterfactual questions such as ‘based on this specific gene expression profile, what would the corresponding chromatin accessibility profile have looked like?’. This could be further combined with in-silico perturbations, i.e., generating synthetic profiles of one modality and using the model to infer corresponding profiles in other modalities. Additionally, this technique can be used to query the trained model for, e.g., subpopulations of cells where the cross-modality predictions are particularly well or particularly poorly aligned with the true measurements and to further characterize them, thus, also facilitating interpretability.

Examples of approaches that employ this technique are given by, e.g., Wu et al. [62], Minoura et al. [45] and Zhao et al. [70], and will be presented in more detail below in Section 3 and in the experimental Section 6

### 2.4 Adversarial training strategies

Another commonly used regularization technique is given by adversarial training, which is closely related to cross-modality translation and is often employed concurrently. Such adversarial components are often integrated into a variational or standard autoencoder framework and are inspired by generative adversarial networks (GANs) [21], another form of DGMs that differs from VAEs in how the joint probability distribution over all input features is specified. While VAEs learn an explicit parameterization of (an approximation of) this distribution (see 2.1), in GANs, this distribution is available only implicitly via sampling. A GAN consists of a generator and a discriminator neural network that can be thought of as playing a zero-sum minimax game: The generator simulates synthetic observations that are presented to the discriminator together with real data observations. The discriminator then has to decide whether a given sample is a real observation or a synthetic one from the generator.

In multi-omics data integration, such adversarial approaches are typically integrated into (V)AE models as additional components to regularize the latent representation and/or the decoder reconstructions [24, 36, 65, 70], while, e.g., Amodio and Krishnaswamy [2], Amodio et al. [3] present purely GAN-based approaches. More specifically, a discriminator is typically employed to distinguish between two omics modalities, either based on samples from their latent representations or based on reconstructed samples from cross-modal decoders (Figure 1 E, black arrows). The objective of the discriminator then is to maximize the probability of correctly identifying the original modality a sample comes from, while the encoder and decoder of the (V)AE model are trained to fool the discriminator by producing samples that are indistinguishable. By training all components jointly, the (V)AE model is encouraged to find a latent embedding in which the different modalities are better aligned and integrated, and/or learn decoders that allow for accurate cross-modal predictions well aligned with the intra-modal predictions. In practice, this is achieved by incorporating adversarial penalty terms into the loss function.

Such adversarial components can also be used to train the model in a cyclical fashion for additional intra-modal and cross-modal consistency. For intra-modal consistency, the low-dimensional embeddings of samples of one modality are decoded with the modality-specific decoder. Subsequently, the reconstructions are re-encoded with the modality-specific encoder and compared to the original embedding of the sample. An adversarial discriminator can be employed to align the embedding of the original sample with the embedding of the re-encoded reconstruction of that sample (Figure 1 E, red arrows). For cross-modal consistency, the low-dimensional embeddings from samples of one modality are decoded and subsequently re-encoded with the decoder and encoder of the other modality. By aligning these cross-modal embeddings with the original embeddings using an adversarial discriminator, the model can learn to produce cross-modal translations that are consistent with the original sample when re-embedded in the latent space.

## 3 Literature review

Although recently, several available deep learning-based applications for the integration of single-cell multi-omics data have been reviewed in [17] and [51], there is still a lack of a more comprehensive review focusing specifically on DGMs. In the following, we are going to survey approaches for paired (both modalities measured in the same cell in one experiments) and unpaired (modalities measured in different cells in separate experiments) single-cell data. An overview is given in Table 1, where we list recent deep learning-based approaches for multi-omics data integration. We remark whether the methods are designed for paired or unpaired datasets and compare the basic network architectures and demonstrated modalities on which the respective methods have been demonstrated. Additionally, we comment on the integration tasks tackled by each model and provide a reference to the implementation.

**Table 1:**
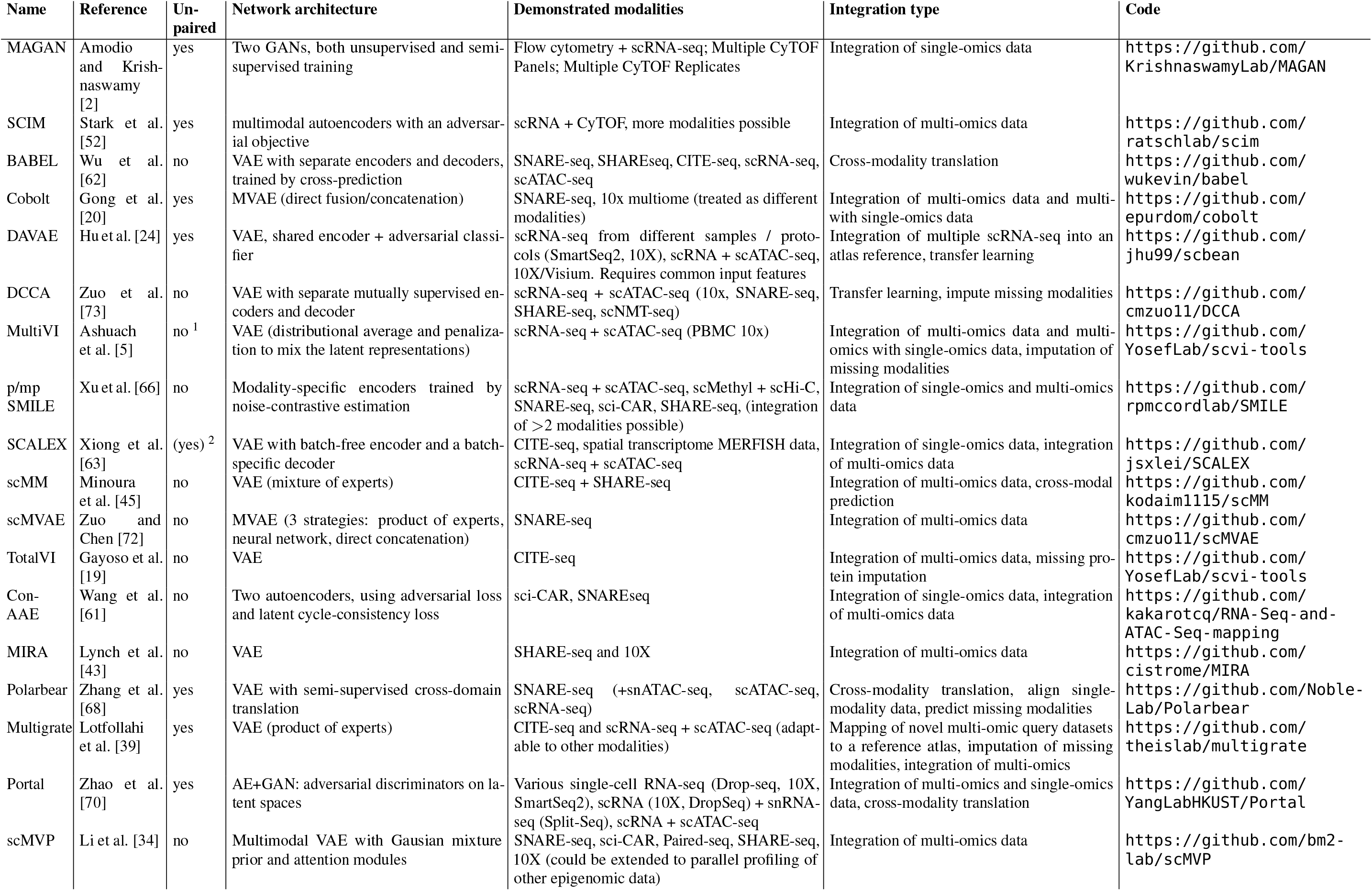
Overview of recently published deep learning-based methods to integrate single-cell multi-omics data. ^x^Only for mapping single-omics to multi-omics; ^2^Only when converting peaks to activity scores.

We exclusively included methods that learn a joint embedding based on DGMs and have been demonstrated on multi-omics data of different modalities (not just, e.g., single-cell RNA-seq from different protocols).

### 3.1 Approaches for paired data

The Cobolt model [20] learns shared representations between modalities and is based on a multimodal VAE, where an independent encoder network is used for each modality and the learned parameters of the posterior distributions are combined using a PoE approach. Additionally, Cobolt can jointly integrate single-modality datasets with multi-omics datasets, allowing one to draw on the many publicly available scRNA-seq or scATAC-seq datasets.

Multigrate [39] is another model that employs a PoE to combine the posteriors of different modalities. Additional datasets can be integrated into the model by minimizing the maximum mean discrepancy (MMD) loss between joint representations of different datasets.

Similar to Cobolt and Multigrate, scMM [45] is a VAE-based method that trains an encoder network for each modality independently. However, instead of combining the parameters of the posterior distributions using a PoE, a MoE is used. By equally mixing information from both modalities through the MoE, the model avoids putting too much emphasis on one individual modality only [45]. In addition, scMM provides a method for model interpretability that uses latent traversals, where synthetic cells are generated by the learned decoder and one latent variable is modified continually, while the others remain fixed. The Spearman correlations calculated between each latent variable and the features of each modality then allow relevant features to be identified. Additionally, by using a Laplace prior, scMM learns disentangled representations, with correlations between latent variables being penalized, which allows for better interpretation of individual features [55].

Similarly, the MultiVI model presented by Ashuach et al. [5] is also based on a MoE with *α_m_* = 1/*M* where *M* denotes the number of modalities, as the authors use individual encoders for each data modality and then average the resulting variational posteriors. However, a regularization term is added to the ELBO, which penalizes the distance between the learned latent representations such that a joint representation can be inferred [5].

While the single-cell multi-view profiler (scMVP) [34] is also based on a multimodal VAE architecture with modality-specific encoders and decoders and a joint latent space, it more explicitly accounts for the much higher sparsity of single-cell measurements from joint profiling protocols, with a throughput of only one-tenth to one-fifth of that of single-modality assays [34]. Specifically, the authors employ attention-based building blocks for both the encoder and decoder. Attention mechanisms have first been proposed in computer science in the context of machine translation [6, 28] and are based on the idea of using flexible weighting of an input observation, to have the model specifically ‘attend to’ the most important parts of the observation. In the context of omics data, attention scores are assigned to the observed features (e.g., genes, chromatin loci) of each cell, to enhance the effect and interplay of specific features. In contrast to fixed weights, the attention scores are learned during model training and can thus adapt to highlight the most informative features for learning, e.g., latent representations. Attention-based mechanisms have specifically been popularized by transformer models [57] due to their high performance on sparse datasets in the area of natural language processing or protein structure prediction. In scMVP, the authors build on that by using multi-head self-attention transformer modules to capture local, long-distance correlation in the encoder and decoder of the term frequency-inverse document frequency-transformed [53] scATAC-seq data while using simple attention blocks in the RNA encoder and decoder. Given the latent embedding, the modality-specific decoders are weighted according to the posterior probabilities of cell-type or cluster identity. To encourage consistency of the shared latent space, the decoder-reconstructed values of each modality are again embedded into the latent space, and the KL-divergence between the joint latent embedding and the modality-specific re-embedding from the reconstructed data is minimized as an additional loss term. This corresponds to the idea of cyclical adversarial training as described in Section2.4 and Figure 1 E. More generally, this concept is based on a cycle GAN [71] and is also present in, e.g., Khan et al. [26], Wang et al. [61], Xu et al. [65], Zhao et al. [70] and Zuo et al. [73].

SCALEX [63] builds on SCALE (Single-Cell ATAC-seq Analysis via Latent feature Extraction) [64], a tool for analyzing scATAC-seq data. The developers of SCALE found that its encoder could be beneficial in disentangling cell-type- and batch-related features, which would allow for online integration of different batches. Specifically, using a VAE, SCALEX integrates different batches into a batch-invariant embedding through simultaneous learning of a batch-free encoder and a batch-specific decoder. The latter contains a domain-specific batch normalization layer. This allows the encoder to concentrate only on batch-invariant biological data components while being oblivious to batch-specific variations. The resulting generalizability of the encoder further allows for the integration of new single-cell data in an online manner, i.e., without the need to retrain the model. The authors demonstrate this property of SCALEX by generating multiple expandable single-cell atlases.

Another subgroup of models addresses the task of translating between different modalities. These cross-modality translation approaches, however, often do not learn a common latent representation of the data. For example, Polarbear [68] trains VAEs on each of two modalities (here: scRNA-seq and scATAC-seq data) and then links the respective encoders to the decoders of the other modality. The authors intend that the training in the first stage, i.e., the training of the individual VAEs, takes place on publicly available single-assay data, whereby the translation task is carried out on SNARE-seq data in a supervised manner.

Another such model called BABEL [62] similarly employs distinct modality-specific encoders and decoders for scRNA- and scATAC-seq data but utilizes a shared latent space. In contrast to PoE/MoE approaches, this joint representation is not constructed from separate spaces from each modality, but the encoders directly project onto the common latent space. Mutual cross-modal translation together with single-modality reconstruction are then used to train the model, i.e., from each modality-specific encoder, a sample of the joint latent representation is obtained and subsequently passed through both decoders to reconstruct both the scRNA and the scATAC profiles of the respective cell. Thus, both the reconstruction of the modality itself and the respective other modality based on the joint latent embedding are evaluated for each modality.

A similar approach is taken by Portal [70], where a domain translation framework is combined with an adversarial training mechanism to integrate scRNA- and scATAC-seq data. Specifically, as in [62], modality-specific encoders directly embed the data in a shared latent space and cross-modal generators are introduced to decode the latent representation to the respective other modality. The resulting domain translation networks for each modality are then trained to compete against adversarial discriminators on the domain of each modality that aims to distinguish between original cells from the respective modality and cells translated from the other modality. The discriminators are specifically designed to adaptively distinguish between domain-shared and domain-unique cells by thresholding the discriminator scores. Since, according to the authors, domain-unique cell populations are prone to be assigned with extreme discriminator scores, discriminators are, thus, made effectively inactive on cells with a high probability of being modality-specific, which avoids the risk of over-correction by enforced alignment of domain-unique cells. Further, additional regularizers are employed: an autoencoder loss based on the within-modality reconstructions, a latent alignment loss to encourage the consistency of a specific cell’s embedding and the embedding of its cross-modal reconstruction, and a cosine similarity loss between cells and their cross-modal reconstructions. Notably, Portal uses the first 30 principal components of a joint PCA as inputs for the model and employs a 20-dimensional latent space, such that the dimension reduction component is less pronounced than for the other models, and the data are not modeled as counts.

The authors of Zuo and Chen [72] have extended scMVAE and proposed Deep Cross-Omics Cycle Attention (DCCA) [73], which improves some of the weaknesses of scMVAE. DCCA combines VAEs with attention transfer. While scMVAE combines two modalities into a shared embedding, which potentially attenuates modality-specific patterns, in the case of DCCA, each data modality is processed by a separate VAE. These VAEs can then learn from each other through mutual supervision based on semantic similarity between the embeddings of each omics modality.

In the sciCAN model presented by Xu et al. [65], modality-specific autoencoders map the input data to a latent space for each modality, and a discriminator is employed to distinguish between the two modalities based on their latent representations. Additionally, a cross-modal generator is employed that generates synthetic scATAC-seq data based on the scRNA-seq latent representation, and a second discriminator is employed to distinguish between generated and real scATAC-seq samples. Additionally, the generated scATAC-seq data can be fed to the encoder again, and the latent representation is compared with the original latent representation from the scRNA-seq data used for generating the scATAC-seq data, thus introducing a cycle consistency loss (see Figure 1 E, Section 2.4). Notably, the model does not necessarily expect paired measurements from the same cell but employs a shared encoder for both modalities, and, thus, requires a common feature set.

The authors of Hu et al. [24] propose the DAVAE model based on domain-adversarial and variational approximation to integrate multiple single-cell datasets and paired scRNA-seq and scATAC-seq data. The model employs an adversarial training strategy to remove batch effects and enable transfer learning between modalities, by incorporating a domain classifier that tries to determine the batch or modality label based on the latent representation of VAE and training the VAE encoder to ‘fool’ the classifier via an adversarial loss component. Similarly to Portal and sciCAN, the DAVAE model also employs a shared encoder and thus requires a common set of input features.

Similarly, the scDEC model proposed by Liu et al. [36] is based a pair of generative adversarial models to learn a latent representation. While focusing on scATAC-seq data analysis, this approach also allows for integrative analysis of multi-modal scATAC and scRNA-seq datasets for trajectory inference during differentiation processes and cell type identification based on the joint latent representation.

Finally, MIRA [43] combines probabilistic cell-level topic modeling [8] with gene-level regulatory potential (RP) modeling [48, 60] to determine key regulators responsible for fate decisions at lineage branch points. The topic model uses a VAE with a Dirichlet prior to learn both the topic of the gene transcription and the topic of gene accessibility for each cell to derive the cell’s identity. Complementing MIRA’s topic model, its RP model integrates the transcription and accessibility information for each gene locus to infer how the expression of the respective gene is influenced by surrounding regulators.

To this end, the topic model learns the rate with which the regulatory influence of enhancers decays with increasing genomic distance. In addition, the identity of key regulators is identified by analyzing transcription factor motif enrichment or occupancy.

### 3.2 Approaches for unpaired data

Since the generation of multi-omics measurements in the same cell is still costly and experimentally complex, many methods for integrating datasets measured in different cells are being developed.

Because of the difficulty of linking latent representations learned from variational autoencoders in the absence of measurement pairing information, Lin et al. [35] proposed a transfer learning approach. Although not a DGM, it is worth mentioning in this article because of its usefulness and the possibility of adapting it to unsupervised settings. Notably, it represents a method for a horizontal alignment task, i.e., it relies on a common set of features as anchors and thus requires the translation of scATAC peaks to gene activity scores.

In a similar spirit, the scDART model proposed by Zhang et al. [69] learns a neural network-based joint embedding or unpaired scRNA-seq and scATAC-seq data by composing the embedding network with a gene-activity module network that maps scATAC peaks to genes. In addition, scDART can leverage partial cell matching information by using it as a prior to inform the training of the gene activity function.

Similar to the sciCAN model presented by Xu et al. [65], scAEGAN [26] also embraces the concept of cycle consistency, integrating the adversarial training mechanism of a cycle GAN [71] into an autoencoder framework. Specifically, for each modality, a discriminator and a generator are defined. In addition to the standard GAN loss for each modality, a cycle loss is calculated by mapping a cell from one modality to the second modality with the second modality’s generator and mapping it back to the first modality with the first modality’s generator and comparing that to the original observation. Unlike for Xu et al. [65], the model does not rely on a common feature set but first trains an autoencoder model independently for each modality before training a cycle GAN on the two latent spaces to enforce their consistency.

A similar approach is employed in the Contrastive Cycle Autoencoder (Con-AAE) proposed by Wang et al. [61]. Again, the consistency between latent spaces of modality-specific autoencoders is enforced by a cycle consistency loss. However, here, it is more tightly integrated within the AE architecture, as the modality-specific encoder and decoders are used as generators, i.e., samples from one modality are embedded with the modality-specific encoder but decoded with the decoder of the other modality, and subsequently encoded with the other modality encoder back to the latent space, where they are compared with the original latent representation from the original encoder of the modality.

A purely GAN-based approach to integrating unpaired data by aligning the respective manifolds is presented in Amodio and Krishnaswamy [2].

Another line of research for the integration of unpaired multi-omics data focuses on the concept of optimal transport [46]. A separate embedding or distance matrix is constructed from each modality, and the alignment task is formulated to find an optimal coupling between the two embeddings or distance matrices. An optimal coupling corresponds to finding a map along which one modality can be “transported” with minimal cost to the other, which can be formalized as an optimal transport problem [46]. Examples for such optimal transport-based methods are UnionCom [10], SCOT [15] and Pamona [11]. While these approaches typically rely on computing a coupling between modality-specific distance matrices and are not deep learning-based, a recent approach called uniPort employs a VAE architecture and solves an optimal transport problem in the latent space. More specifically, a shared encoder that requires a common input feature set across modalities is used to project the data into a common latent space, is combined with modality-specific decoders for reconstruction, and an optimal transport loss is minimized between the latent cell embeddings from different modalities.

Finally, the recently published Graph-Linked Unified Embedding (GLUE) framework [12] is based on the construction of a guidance graph based on prior knowledge of the relations between features of the different modalities to explicitly model regulatory interactions across different modalities with distinct feature spaces. This is achieved by learning joint feature embeddings from the knowledge graph with a graph VAE and linking them to modality-specific autoencoders. Specifically, the decoder of these modality-specific AEs is given by the inner product of the feature embeddings and the cell embeddings from the latent space of the respective modality. Additionally, the cell embeddings of different modalities are aligned using an adversarial discriminator.

## 4 Benchmark Dataset

To acquire an objective performance estimate of the ability of different multi-omics integration approaches to describe the biological state of a cell through learning a joint embedding from multiple modalities, we used the benchmark dataset which was provided in the course of the NeurIPS 2021 competition and for which the ground-truth cell identity labels are known [40]. This dataset was the first available multi-omics benchmarking dataset for single-cell biology. It mimics realistic challenges researchers are faced with when integrating single-cell multi-omics data, e.g., by incorporating nested donor and site batch effects [33].

Specifically, the NeurIPS benchmark dataset is a multi-donor (10 donors), multi-site (4 sites), multi-omics bone marrow dataset comprising two data types [33]:

- CITE-seq data with 81,241 cells, where for each cell RNA gene expression (GEX) and cell surface protein markers using antibody-derived tags (ADT) are jointly captured.
- 10X Multiome assay data with 62,501 cells, where nucleus GEX and chromatin accessibility measured by assay for transposase-accessible chromatin (ATAC) are jointly captured.

In total, this dataset contained information on the accessibility of 119,254 genomic regions, the expression of 15,189 genes, and the abundance of 134 surface proteins, and has been preprocessed as described in Luecken et al. [40]. We acquired the benchmark dataset from the NeurIPS 2021 website (https://openproblems.bio/neurips_2021), it can, however, also be accessed via https://www.ncbi.nlm.nih.gov/geo/query/acc.cgi?acc=GSE194122.

As recommended in Luecken and Theis [42], we filtered this dataset for highly variable genes, as they are considered to be most informative of the variability in the data. In addition, analogous to the FindTopFeatures function of Signac [53], we filtered the ATAC data such that we retained only peaks with the 25 % highest overall counts. Finally, to determine the effect of the number of cells, we randomly subsampled the original NeurIPS dataset to subsamples containing information on 500, 1000, 2500, 5000, and 10 000 cells, where for each number of cells we sampled 10 subsamples of that size.

## 5 Performance metrics

Generating a highly resolved, interpretable, low-dimensional embedding capturing the underlying biological cell states is pivotal for the analysis of multi-omics data [32, 33]. We assess the performance of the compared integration approaches based on six metrics capturing the conservation of biological variation (normalized mutual information (NMI), cell type average silhouette width (ASW), trajectory conservation) and the degree of batch removal (batch ASW, site ASW, graph connectivity) [33]. These metrics are described in detail in Luecken et al. [41] and are briefly introduced below:

- **NMI** compares the overlap of two clusterings. It is used to compare the Louvain clustering of the joint embedding to the cell type labels. It ranges from 0 (uncorrelated clustering) to 1 (perfect match).
- **Cell type ASW** is used to evaluate the compactness of cell types in the joint embedding. It is based on the silhouette width, which measures the compactness of observations with the same labels. Here, the ASW was computed on cell identity labels and scaled to a value between 0 (strong misclassification) and 1 (dense and well-separated clusters).
- The **trajectory conservation** assesses the conservation of a continuous biological signal in the joint embedding. Trajectories computed using diffusion pseudotime after integration for relevant cell types are compared. Based on a diffusion map space embedding of the data, an ordering of cells in this space can be derived. Using Spearman’s rank correlation coefficient between the pseudotime values before and after integration, the conservation of the trajectory can be quantified, with the scaled score ranging from 0 (reverse order of cells on the trajectory before and after integration) to 1 (same order).
- **Batch ASW** describes the ASW of batch labels per cell. The scaled score ranges from 0 to 1, where 1 indicates well-mixed batches and any deviation from 1 indicates a batch effect.
- **Site ASW** describes the ASW of site labels per cell and can be interpreted analogously to batch ASW.
- The **graph connectivity** score evaluates whether cells of the same type from different batches are close to each other in the embedding by assessing if they are all connected in this embedding’s k-nearest neighbor (kNN) graph. It ranges from 0 (no cell is connected) and 1 (all cells with the same cell identity are connected).

## 6 Results

We use various metrics to quantify the preservation of biological variation and metrics for the removal of technical effects based on the 10-dimensional embeddings obtained when applying Cobolt, scMM, TotalVI, and SCALEX to subsamples of the NeurIPS CITE-seq dataset, and Cobolt, scMM, MultiVI, scMVP, DAVAE, and Portal to subsamples of the NeurIPS Multiome dataset. We randomly sampled 500, 1000, 2500, 5000, and 10000 cells ten times each and applied the models to the respective datasets. We refrain from extensive parameter optimisation as we put ourselves in the position of a user new to the field of deep learning, who will, most likely, leave the default parameters unchanged and use the same parameters as the original authors in their application of their proposed method. Thus, we used the default hyperparameters of the respective models as reported by the authors who originally proposed them where possible (Supplementary Material: Hyperparameters).

When applying scMM to the CITE-seq data, we frequently observed non-converging training runs, in particular for larger sample sizes. Here, we refer to the convergence of the iterative optimization procedure by stochastic gradient descent on the loss function of the respective model (see also Section 2). Convergence is achieved if towards the end of the training, the changes in the loss function in each iteration become smaller and eventually level out, whereas in non-converging runs we observe exploding gradients of the loss function. This is often due to suboptimal hyperparameter choices. For scMM, lowering the learning rate for sample sizes above 2500 by one order of magnitude and increasing the batch size from 128 (default used by scMM) to 200 achieved convergence of the model training on all subsamples.

In general, similar performances were achieved irrespective of which of the two data types we used for deriving a joint embedding (**Figures 4 and 5)**. For the Multiome dataset, two of the considered tools, DAVAE and Portal, employ a shared encoder based on a common set of features across both modalities (top 30 principal components of a joint PCA on both datasets for Portal and common highly variable genes when converting scATAC peaks to gene activity scores for DAVAE) and thus embed each cell’s profiles in the two modalities separately. To keep the evaluation as comparable as possible to the other tools, we thus created a joint embedding by calculating the mean of each cell’s embedded profiles in the two modalities in a mixture-of-experts approach.

We compare our results with the metric values achieved by the models of the NeurIPS 2021 competition for the integration of the Multiome dataset (data points were extracted via WebPlotDigitizer-4.5 [49] from Supplementary Figure 6 of [33]). However, as we merely used a subset of at most 10,000 cells of the original benchmark dataset, we expect our investigated algorithms to score higher for most metrics if they were to be subjected to the complete benchmark dataset.

By visual inspection of the Uniform Manifold Approximation and Projection (UMAP) (Becht et al. [7], Konopka and Konopka [31] version 0.2.9.0 with default parameters) plots of one exemplary subsample (**Figure 2** and **Figure 3**), we see that MultiVI shows no obvious clustering for 500 cells (2, top panel). In contrast, defined cluster structures are beginning to build at this low cell number, and become more refined for 10,000 cells, for all other investigated tools. This behavior of MultiVI for smaller numbers of cells is also reflected in lower values for most of the investigated performance metrics (**Figures 2 and 3)**. Interestingly, the TotalVI tool, which is built on a similar architecture and was used for the CITE-seq dataset does not show such behavior (4, top panel).

**Figure 2:**
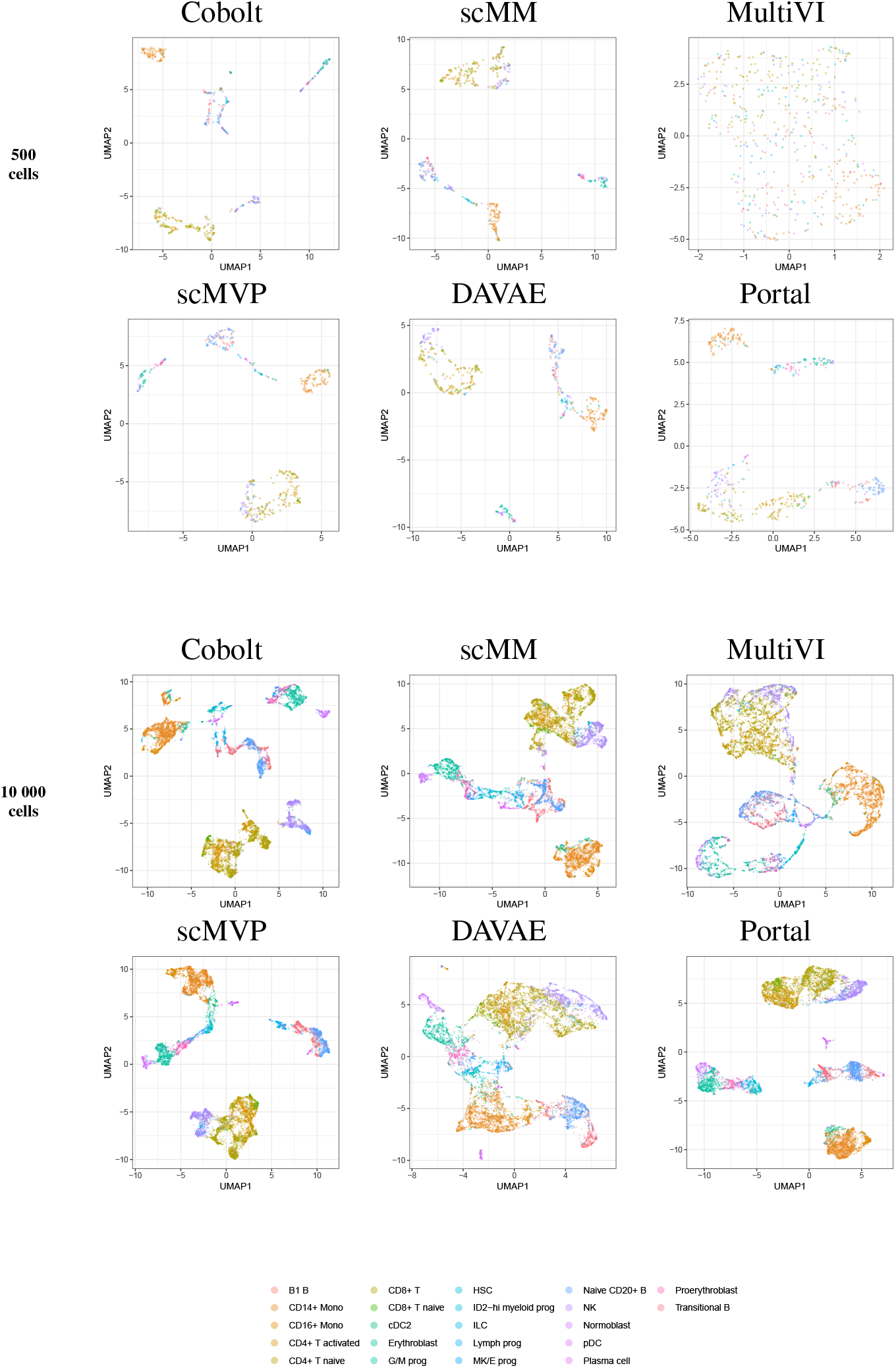
UMAP of the 10-dimensional latent space of Cobolt, scMM, MultiVI, scMVP, DAVAE, and Portal based on 500 (top) and 10000 (bottom) cells of one exemplary subsample from the Multiome dataset each. The color coding corresponds to manually annotated cell types as provided by Luecken etal. [40].

**Figure 3:**
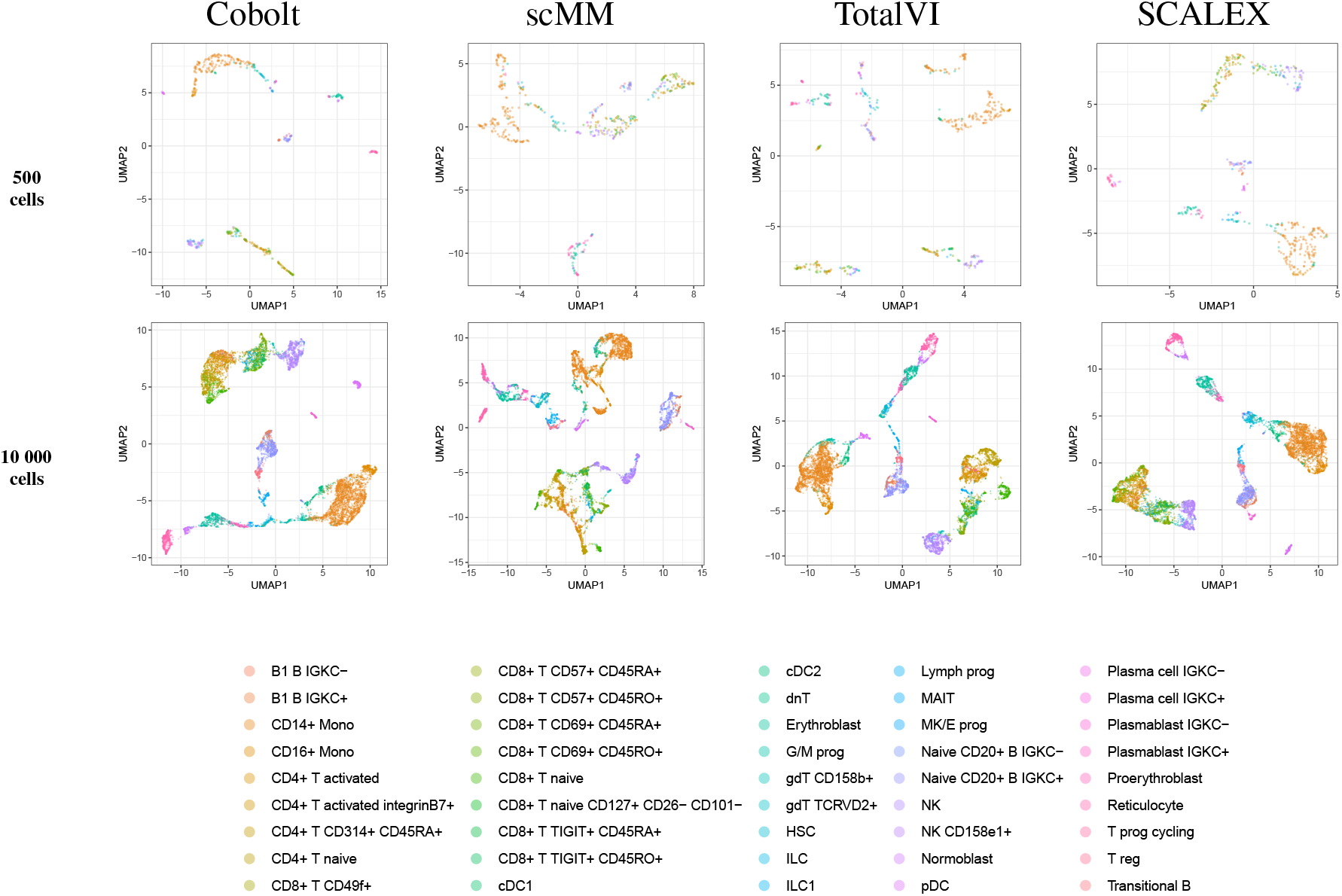
UMAP of the 10-dimensional latent space of Cobolt, scMM, TotalVI, and SCALEX based on 500 (top) and 10000 (bottom) cells of one exemplary subsample from the CITE-seq dataset each. The color coding corresponds to manually annotated cell types as provided by Luecken et al. [40]. The following cell types are not present in the 500 cell sample: CD4+ T CD314+ CD45RA+, CD8+ T naive CD127+ CD26-CD101-, cDC1, dnT, Plasma cell IGKC-, Plasma cell IGKC+, Plasmablast IGKC-, T prog cycling.

**Figure 4:**
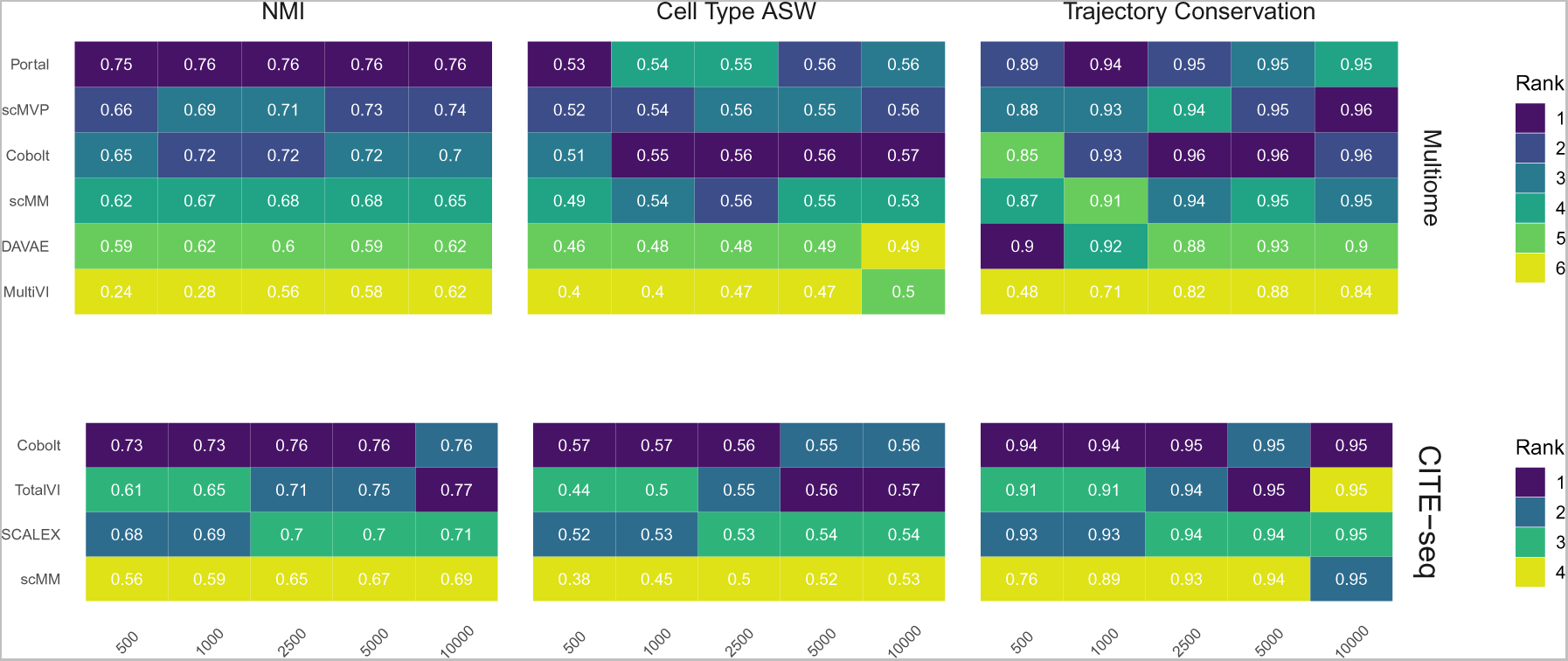
Biological preservation metrics. NMI, cell type ASW, and trajectory conservation score indicate the biological preservation quality achieved by joint embeddings from various models. On the 10x Multiome data, the performance of Portal, scMVP, Cobolt, scMM, DAVAE, MultiVI is shown (top), whereas the performance on CITE-seq data is shown for Cobolt, TotalVI, SCALEX, and scMM (bottom). Median scores across all iterations are shown inside the tiles.

**Figure 5:**
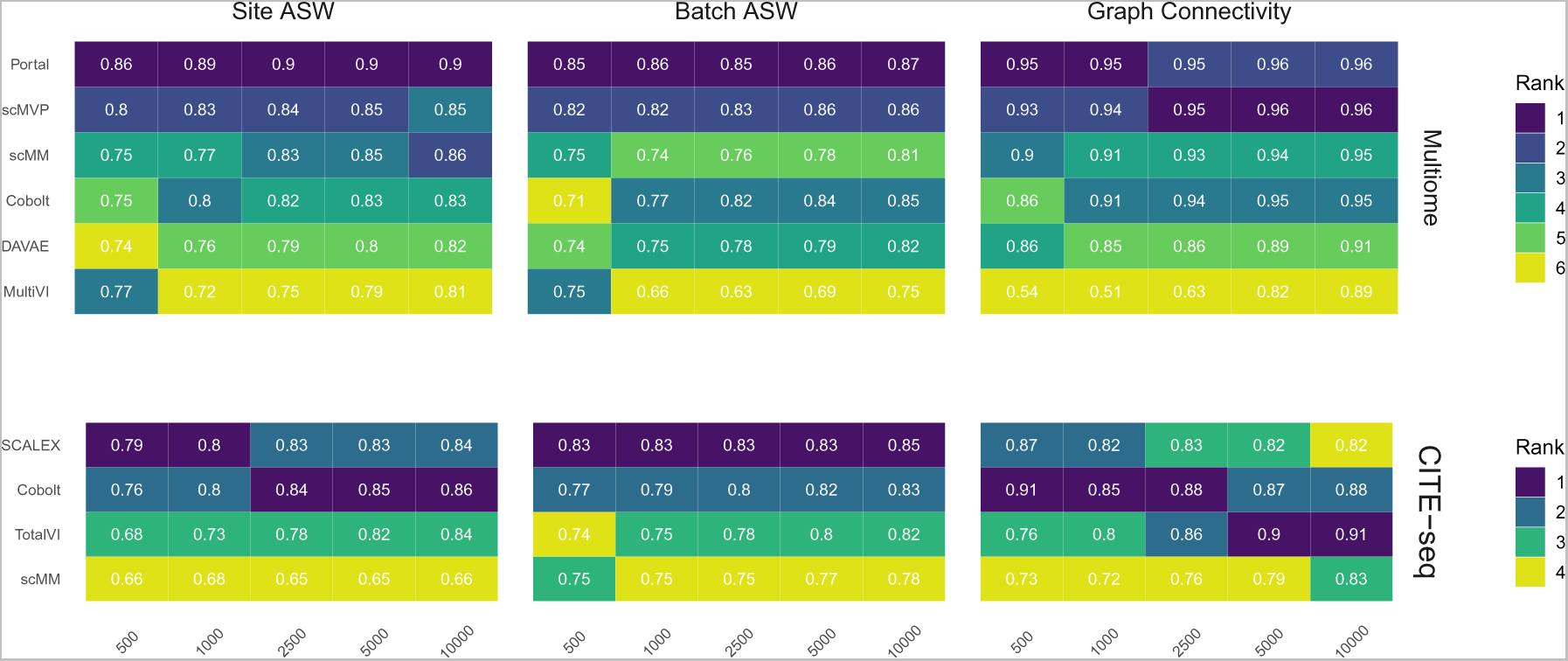
Technical effect removal metrics. Site ASW, batch ASW, and graph connectivity score indicate the quality of technical effects removal achieved by joint embeddings from various models. On the 10x Multiome data, the performance of Portal, scMVP, scMM, Cobolt, DAVAE, and MultiVI is shown (top), whereas the performance on CITE-seq data is shown for SCALEX, Cobolt, TotalVI, and scMM (bottom). Median scores across all iterations are shown inside the tiles.

UMAP plots including further meta information on the embedded cells are given in **Supplementary Figures 3 to 22** for the exemplary subsample.

To ensure that the number of parameters in the respective models is not the determining factor for decreasing performance on small sample sizes, we calculated the Spearman correlation coefficient between the ranks of the models from **Figures 4 and 5** and the evaluation metrics. The predominantly negative correlations, i.e., lower rank (better performance) with an increasing number of trainable parameters, indicate that more complex models also deliver better performance regardless of the number of observations.

### 6.1 Preserving biological information

We assess the preservation of biological variation based on the NMI, cell type ASW, and the trajectory conservation scores (**Figure 4**). In addition, we show boxplots of the metrics for all models and sample sizes for both Multiome and CITE-seq data in the **Supplementary Figures 1** and **2**, to show the variability of each metric across the 10 replicates of each dataset size.

NMI, as a measure of cluster overlap, reaches values of approx. 0.7 for all Multiome and CITE-seq integrating models. The NMI is slightly lower than what was achieved during the NeurIPS 2021 competition, where the best competition entries reached an NMI of close to 0.8 for the complete Multiome dataset [33] (see **Supplementary Figure 1**). This is to be expected as we evaluate the models in a low sample size scenario. MultiVI profits greatly from a larger cell number, while an increasing cell number only slightly increases the performance of the other models. Across most sample sizes, Cobolt performed best for the CITE-seq datasets, while Portal performs best on the Multiome datasets for all sample sizes but does not profit much from increasing sample size. For larger sample sizes, scMVP shows only slightly worse performance than Portal on the Multiome dataset.

Cell type ASW is a measure of cluster compactness and overlap. We see values of around 0.5 for the Multiome and CITE-seq datasets, which implies overlapping of clusters and only a moderate separation. This is slightly lower than the 0.6 that models have achieved in the NeurIPS 2021 competition [33]. For Multiome data, the impact of cell numbers was minor in Cobolt, scMM, Portal, DAVAE and scMVP representations and higher for MultiVI. For CITE-seq data, only scMM and TotalVI show a dependence between cell type ASW and cell number. As expected, increasing the number of cells leads to a decrease in variance.

The trajectory conversation score measures the preservation of a biological signal, e.g. in the form of developmental processes. For the CITE-seq dataset, all models reach comparable scores of around 0.9 irrespective of the cell numbers, with a substantial decrease in variance for larger cell numbers. In contrast, for the Multiome dataset, an increase in cell numbers affects the trajectory conservation score for all models except DAVAE. In particular MultiVI shows a large improvement in performance with increasing cell numbers, while for Portal, scMVP, Cobolt and scMM, the scores increase from around 0.87 to around 0.96. Cobolt performs best for higher cell numbers, while the performance of Portal and scMVP is on par and slightly better than Cobolt for lower cell numbers. The maximum score that models reach in our analysis slightly exceeds the median of the trajectory conservation scores of around 0.9 achieved by models of the NeurIPS 2021 competition [33].

Taken together, Cobolt is the strongest performing model based on almost all biology preservation metrics on the CITE-seq data and regarding cell type ASW on the Multiome data, performing well even in scenarios with small sample sizes. Portal is the strongest performing model on the Multiome data based on NMI and trajectory conservation and performs well on cell type ASW, also showing consistently high performance across sample sizes.

### 6.2 Removing technical effects

We assess the removal of technical artifacts based on the batch ASW and graph connectivity score (**Figure 5**). As a measure of between-site technical variation and to account for the shortcomings of batch ASW (which does not sufficiently account for the nested batch effects of donors and sites) and graph connectivity (which is not sufficiently challenging) [33], we also assess batch ASW with the site as a covariate (‘Site ASW’), as has been suggested by Lance et al. [33].

The Batch ASW score of around 0.8 that we observe in our results indicates only a minor batch effect, although the score is slightly lower than the 0.9 that models achieved in the course of the NeurIPS 2021 competition [33] (see **Supplementary Figure 2**). There is a slight increase in performance for increasing cell numbers across both datasets. For the Multiome dataset, Portal consistently performed best, closely followed by scMVP in particular for larger cell numbers, while MultiVI scored lowest for most cell number settings. For the CITE-seq dataset, SCALEX shows the highest Batch ASW score across all cell number settings, implying superior handling of batch effects even with small sample sizes. This is in line with SCALEX being specifically designed to separate batch-related from batch-invariant components [63].

The graph connectivity score indicates how well cells of the same cell type and cells coming from different batches are connected in the joint embedding. For the Multiome dataset, MultiVI’s graph connectivity score is considerably lower for small sample sizes, while all models improve performance with an increasing number of cells. Portal and scMVP are the best performing models, reaching a score of almost 1 for higher cell numbers in the case of the Multiome dataset in line with the scores achieved by the models of the NeurIPS competition [33]. For the CITE-seq dataset, the performance of TotalVI increased with increasing cell numbers, achieving the highest graph connectivity score for 5,000 and 10,000 cells. In contrast, the number of cells had only a minor effect on the other models. scMM consistently had the lowest graph connectivity score for the CITE-seq dataset.

Site ASW captures site-specific batch effects. Compared to Batch ASW, the performance differences between the models that we applied to the CITE-seq dataset are enhanced. For the CITE-seq dataset, Cobolt and SCALEX perform best, with Cobolt surpassing SCALEX for increasing cell numbers. scMM consistently has the lowest scores on the CITE-seq data. For the Multiome data, the spread of the investigated models is comparable to the one of Batch ASW. Portal achieved the highest Site ASW scores followed by scMVP, which is in agreement with their high Batch ASW score.

Portal and scMVP are the best performing models for metrics considering the removal of technical effects on the Multiome data, whereas MultiVI’s performance suffers. On the CITE-seq data, SCALEX and Cobolt are among the best performing models, while scMM shows consistently low scores across metrics and cell numbers.

### 6.3 Usability

The scMM model by [45] was easily usable. The authors provide both a command line interface and a script that is straightforward to adapt and run. However, HDF5-based data (such as the popular ‘AnnData’ objects) has to be manually restructured to separate files to be used as input for the model. For CITE-seq-data, model training did not always converge, in particular for larger sample sizes, which could be addressed by lowering the learning rate and changing the batch size. While this behavior did not occur with very small learning rates (2 orders of magnitude smaller than the default used by Minoura et al. [45]), this also tended to substantially lower the performance.

To run the scMVP model by [34], package dependency issues had to be resolved manually. Here, too, data had to be restructured manually to fit the custom input data structs defined by the authors. Adapting and running the model and extracting the learned embedding was straightforward.

All in all, all investigated tools were relatively easy to use and adapt, though in most cases not without at least intermediate programming skills (e.g., to transform own data into rather specific and often largely undocumented data structs defined by the authors).

Finally, looking at the time the tools need for their calculations, we found that the central processing unit (CPU) time (without preprocessing) of Cobolt considerably exceeds the the CPU time of the other tools especially for the Multiome dataset (**Supplementary Figures 23 and 24**). Of note, the tools were run on different machines, which hinders a direct comparison. However, it should give the reader a rough idea about the processing time each tools requires, and it is useful to see how well the different investigated tools scale timewise for increasing cell numbers.

## 7 Outlook and Discussion

The rapid emergence of experimental protocols for profiling several omics layers from the same cell or in independent experiments is closely followed by the development of corresponding computational models for analyzing and integrating such data. These methods promise to answer biological questions previously out of reach. Still, they have so far been hampered by often rather small and sparse datasets and the lack of a systematic overview and comparison. In particular, considering the sparsity and high dimensionality inherent to single-cell (multi-)omics data, researchers seek to identify a lowdimensional embedding that integrates the information from multiple modalities and can be used for further downstream analyses. Consequently, many computational tools to infer such a joint latent representation have recently been proposed, often based on deep learning approaches due to their success in identifying complex structures from data in unsupervised settings. Specifically, deep generative models such as VAEs that infer a low-dimensional, compressed representation of the input data in an unsupervised way are among the most popular solutions, often including additional components or custom architectures to accommodate the properties of single-cell multi-omics data and facilitate specific characteristics of the learned embedding.

Due to the rapidly growing number of complex methodological proposals for solving the challenging task of computationally integrating multi-omics data, an overview and categorization of such models are essential for understanding the advantages and disadvantages of the different methods. We have compiled a comprehensive review of the literature on DGMs for learning joint embeddings of multi-omics data and categorized the different models according to their architectural choices.

In addition to this overview, we have also illustrated the robustness of selected models to small sample sizes, where sample size refers to the number of cells in the dataset. For evaluating model performance, we have relied on the guidelines of a comprehensive benchmarking project [40]. We have evaluated the models based on established metrics concerning their ability to adjust for technical effects while maintaining biological signals. Our analyses have shown that Cobolt, an approach that uses a multimodal VAE with products of experts to combine individual embeddings, and Portal, an approach that uses the principal components of a joint PCA on both modalities as input to an autoencoder with an adversarial training strategy, deliver the best performance for most biological preservation metrics, particularly for small numbers of cells. On the other hand, Portal and scMVP, an approach that employs attention-based components and a dedicated architecture to deal with the sparsity of scATAC-seq data, score highest for metrics related to removing technical artifacts on the 10x Multiome data, while SCALEX performs best on the CITE-seq data.

To consider the usability of the approaches from the perspective of a user who is not an expert in tuning deep learning models, we employed the default hyperparameters of the models as proposed by their original authors. While this could potentially introduce bias and dedicated tuning of hyperparameters might improve the results, our focus was on comparing the different approaches relative to each other and relative to the sample size of the respective dataset rather than absolute values of a metric which might be improved by hyperparameter tuning.

Especially for users with little programming experience, some of the models investigated will be difficult to apply, as they require, e.g., the use of command line tools. Here, libraries such as scvi-tools [18] offer a significant benefit by providing extensive documentation and exemplary applications.

Interpretability is an aspect that is of great importance for the application of DGMs [55]. Some of the models we have reviewed already offer the possibility of making the corresponding outputs interpretable for users. For example, post-hoc methods such as applying archetypal analysis [14] to the joint embedding as conducted by TotalVI [19], can make the models explainable after they have been trained. On the other hand, model-based interpretability can be directly incorporated into the model architecture to allow for immediate interpretation, such as the latent traversals and specification of a dedicated prior to facilitate disentanglement in [45]. However, no dominant approach has yet emerged in this area, providing scope for new developments.

We would like to stress that our review should not be understood as a comprehensive benchmark but rather as an illustrative case study, as we merely looked at the investigated DGM tools in the scope of representative examples of the landscape of state-of-the-art approaches, with a focus on potential differences in the number of cells they require to perform well.

In this work, we merely discussed some of all available omics modalities, and the performance of the models may be affected for the better or the worse if applied to other data types due to differing data characteristics, e.g., in the degree of sparsity.

The performances we obtained by running the investigated tools on a benchmark dataset may well deviate if applying those tools to other datasets of differing biological backgrounds, e.g., in terms of cell type composition, tissue types, etc. Although a focus on specific cell types is beyond the scope of our review, we invite others to use our findings as a stepping stone to explore the performance of DGMs for specific biological scenarios.

In the future, linking information from measurements of transcriptomes, epigenomes, proteomes, chromatin organization, etc., could lead to a deeper understanding of cellular processes. Scientists could then further enhance their understanding of these processes by information on the spatial context.

## Author Contributions

E.B., M.H., and M.T. conceived the idea for the manuscript, conducted the analyses, and wrote the manuscript. C.K. and H.B. contributed to writing and proofread the manuscript. All authors read and approved the final manuscript.

## Supplemental Data

### Data Availability Statement

The benchmark dataset can be accessed via https://www.ncbi.nlm.nih.gov/geo/query/acc.cgi?acc=GSE194122.

The joint embeddings learned by all models can be found at https://doi.org/10.5281/zenodo.6616542.

The code to reproduce all analyses and figures can be found at https://github.com/MTreppner/multiomics_dgms.

