## Supplementary Figures and Tables for "The performance of deep generative models for learning joint embeddings of single-cell multi-omics data"

### Supplementary Material

September 15, 2022

#### 1 Hyperparameters

**Cobolt:**

*Multiome data: scRNA-seq + scATAC-seq*
*CITE-seq data: scRNA-seq + surface proteins*
*Version: Status as of 01/02/2022*

- Learning rate for the Adam optimizer [lr]: 0.005
- Number of annealing epochs for the cost annealing scheme [annealing_epochs]: 30
- Parameter of the Dirichlet prior distribution [alpha]: None
- A list of integers indicating the number of hidden dimensions to use for the encoder neural networks [hidden_dims]: None
- Whether to use the intercept term for batch correction [intercept_adj]: True
- Whether to use the slope term for batch correction [slope_adj]: True
- The proportion of random samples to use for training [train_prop]: 1
- Number of epochs/iterations [num_epochs]: 100

**SCALEX:**

*CITE-seq data: scRNA-seq + surface proteins*
*Version: Development version (`https://github.com/ericli0419/SCALEX`) (as suggested by the developers) as of 04/03/2022*

- Filtered out cells that are detected in less than min_features [min_features]: 1 (as suggested by the developers, non-default)
- Filtered out genes that are detected in less than min_cells [min_cells]: 1 (as suggested by the developers, non-default)
- processed: True (as suggested by the developers, non-default)
- Specify the single-cell profile, RNA or ATAC [profile]: 'RNA'

- Use intersection ('inner') or union ('outer') of variables of different batches [join]: 'inner'
- Number of highly-variable genes to keep [n_top_features]: 2000
- Number of samples per batch to load [batch_size]: 64
- Learning rate[lr]: 2e-4
- Max iterations for training. Training one batch_size samples is one iteration [max_iteration]=30000
- Use for new dataset projection. Input the folder containing the pre-trained model. If None, don't do projection. [projection]: None
- Use with projection. If False, concatenate the reference and projection datasets for downstream analysis. If True, only use projection datasets.[repeat]: False
- If True, calculate the imputed gene expression and store it at adata.layers['impute'].[impute]: None
- Number of samples from the same batch to transform [chunk_size]: 20000

**scMM:**
*Version: Status of* `https://github.com/kodaim1115/scMM`*) as of 01/02/2022*
*Multiome data: scRNA-seq + scATAC-seq*

- optimizer: ADAM
- batch size: 64
- learning rate: 1e-4
- number of epochs: 100
- number of warm-up epochs: 50
- latent dimension: 10
- number of hidden layers in encoder and decoder: 3
- dimension of hidden layers for the RNA encoder and decoder: 500
- dimension of hidden layers for the ATAC encoder and decoder: 100

*CITE-seq data: scRNA-seq + surface proteins*

- optimizer: ADAM
- batch size: 200
- learning rate: 2e-3
- number of epochs: 50

- number of warm-up epochs: 25
- latent dimension: 10
- number of hidden layers in encoder and decoder: 3
- dimension of hidden layers for the RNA encoder and decoder: 200
- dimension of hidden layers for the ATAC encoder and decoder: 200

**scMVP:**
*Version: Status of* `https://github.com/bm2-lab/scMVP`*) as of 15/02/2022*

- optimizer: ADAM
- batch size: 128
- learning rate: 5e-4
- number of epochs: 30
- latent dimension: 10
- architecture of RNA encoder: 128-dimensional hidden layer, layer normalization layer, batch normalization layer, relu activation weighted by mask attention
- architecture of ATAC encoder: 128-dimensional hidden layer, batch normalization layer, relu activation, multi-heads self-attention layer with 8 attention heads of dimension 16 followed by layer normalization
- combination of RNA + ATAC encoder with a 256-dimensional linear layer, followed by 128-dimensional layer to produce latent mean and variance
- decoder architecture: like encoder plus additional attention module based on clustered cell types
- cluster number: 10
- weight of KL-divergence in loss: 1.0

**MultiVI:**
*Multiome data: scRNA-seq + scATAC-seq*
*Version: scvi 0.14.5*

- n_latent=10
- max_epochs=100
- lr=0.0001
- train_size=0.9
- validation_size=None

- batch_size=128
- weight_decay=0.001
- eps=1e-08
- early_stopping=True
- save_best=True
- check_val_every_n_epoch=None
- n_steps_kl_warmup=None
- n_epochs_kl_warmup=50
- adversarial_mixing=True
- n_layers_encoder=2
- n_layers_decoder=2
- dropout_rate=0.1
- region_factors=True
- gene_likelihood='zinb'
- use_batch_norm='none'
- use_layer_norm='both'
- latent_distribution='normal'
- deeply_inject_covariates=False
- encode_covariates=False
- fully_paired=False

**TotalVI:**
*CITE-seq data: scRNA-seq + surface proteins*
*Version: scvi 0.14.5*

- n_latent=10
- max_epochs=100
- lr=0.004
- train_size=0.9
- validation_size=None

- batch_size=256
- early_stopping=True
- check_val_every_n_epoch=None
- reduce_lr_on_plateau=True
- n_steps_kl_warmup=None
- n_epochs_kl_warmup=None
- adversarial_classifier=None
- gene_dispersion='gene'
- protein_dispersion='protein'
- gene_likelihood='nb'
- latent_distribution='normal'
- empirical_protein_background_prior=None
- override_missing_proteins=False

### 2 Supplementary Tables and Figures

#### 2.1 Figures

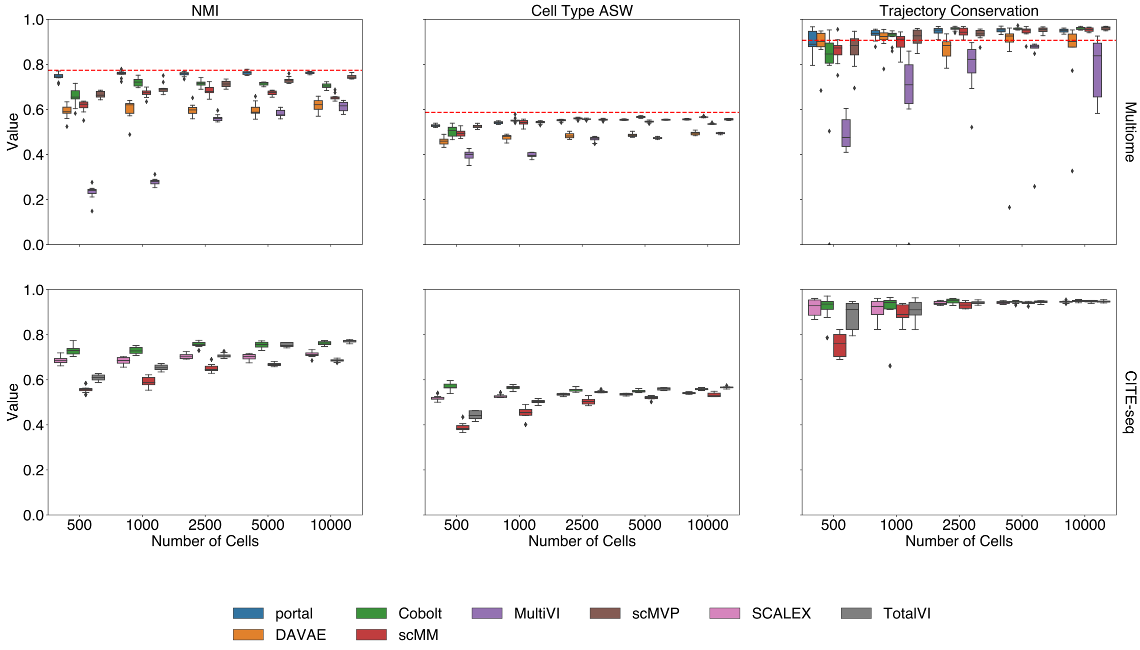


Figure 1: NMI, cell type ASW, and trajectory conservation score indicate the biological preservation quality achieved by joint embeddings from various models. On the 10x Multiome data, the performance of Portal, scMVP, Cobolt, scMM, DAVAE, MultiVI is shown (top), whereas the performance on CITE-seq data is shown for Cobolt, TotalVI, SCALEX, and scMM (bottom). The red dashed lines represent the median performance across all models (excluding the ones with a metric equal to zero) presented and available in the NeurIPS 2021 Multimodal data integration competition [1].

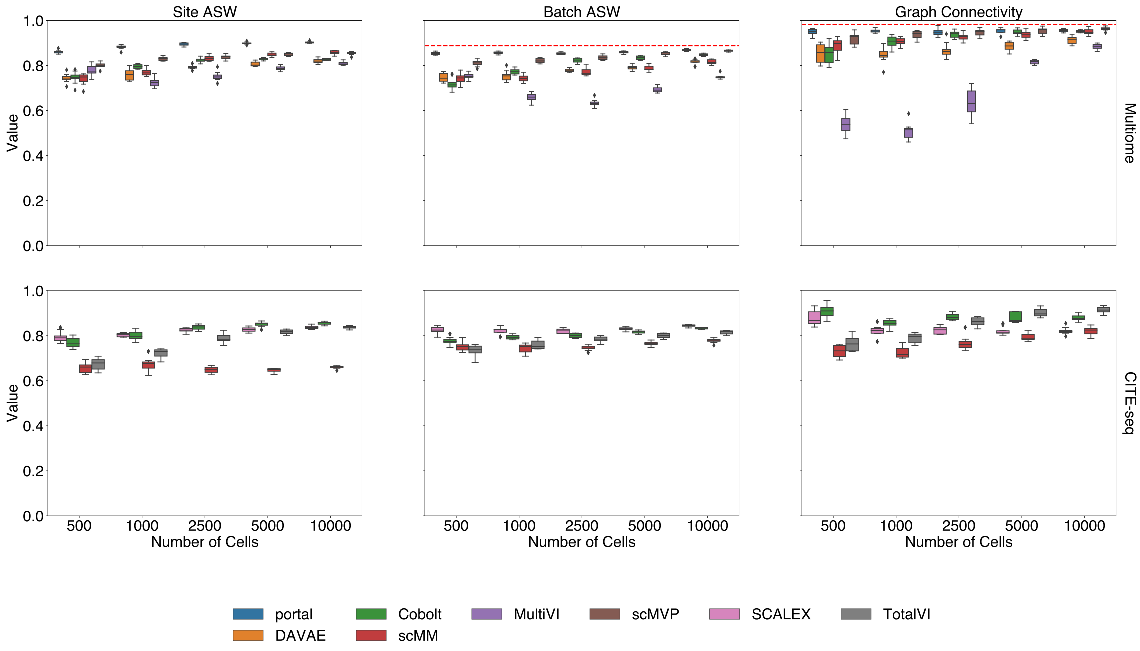


Figure 2: Site ASW, batch ASW, and graph connectivity score indicate the quality of technical effects removal achieved by joint embeddings from various models. On the 10x Multiome data, the performance of Portal, scMVP, scMM, Cobolt, DAVAE, and MultiVI is shown (top), whereas the performance on CITE-seq data is shown for the SCALEX, Cobolt, TotalVI, and scMM (bottom). The red dashed lines represent the median performance across all models (excluding the ones with a metric equal to zero) presented and available in the NeurIPS 2021 Multimodal data integration competition [1].

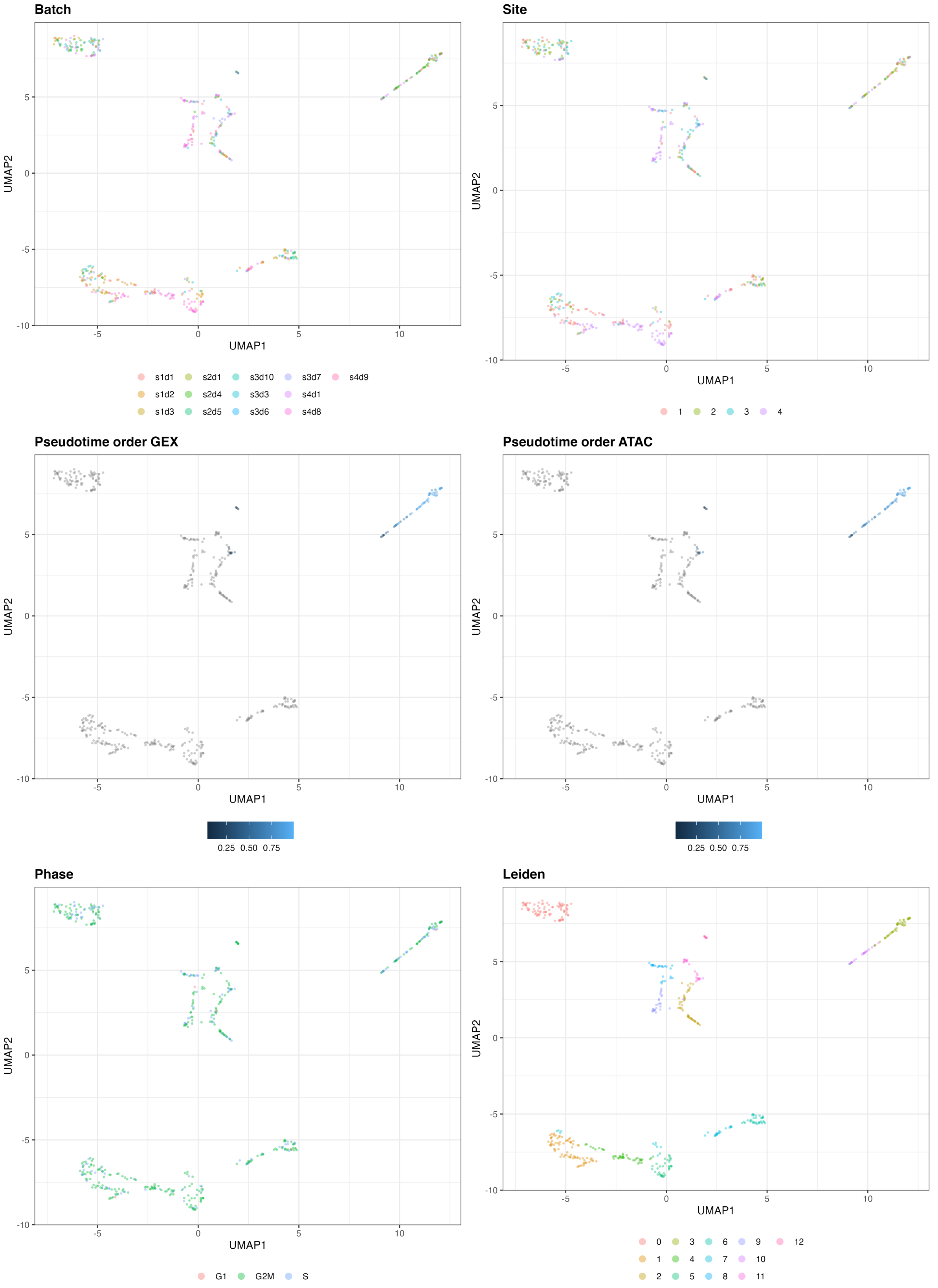


Figure 3: UMAP of the 10-dimensional latent space of Cobolt based on 500 cells of one exemplary subsample from the Multiome dataset. The cells are color-coded by batch, site, pseudotime order GEX, pseudotime order ATAC, phase, and Leiden clustering.

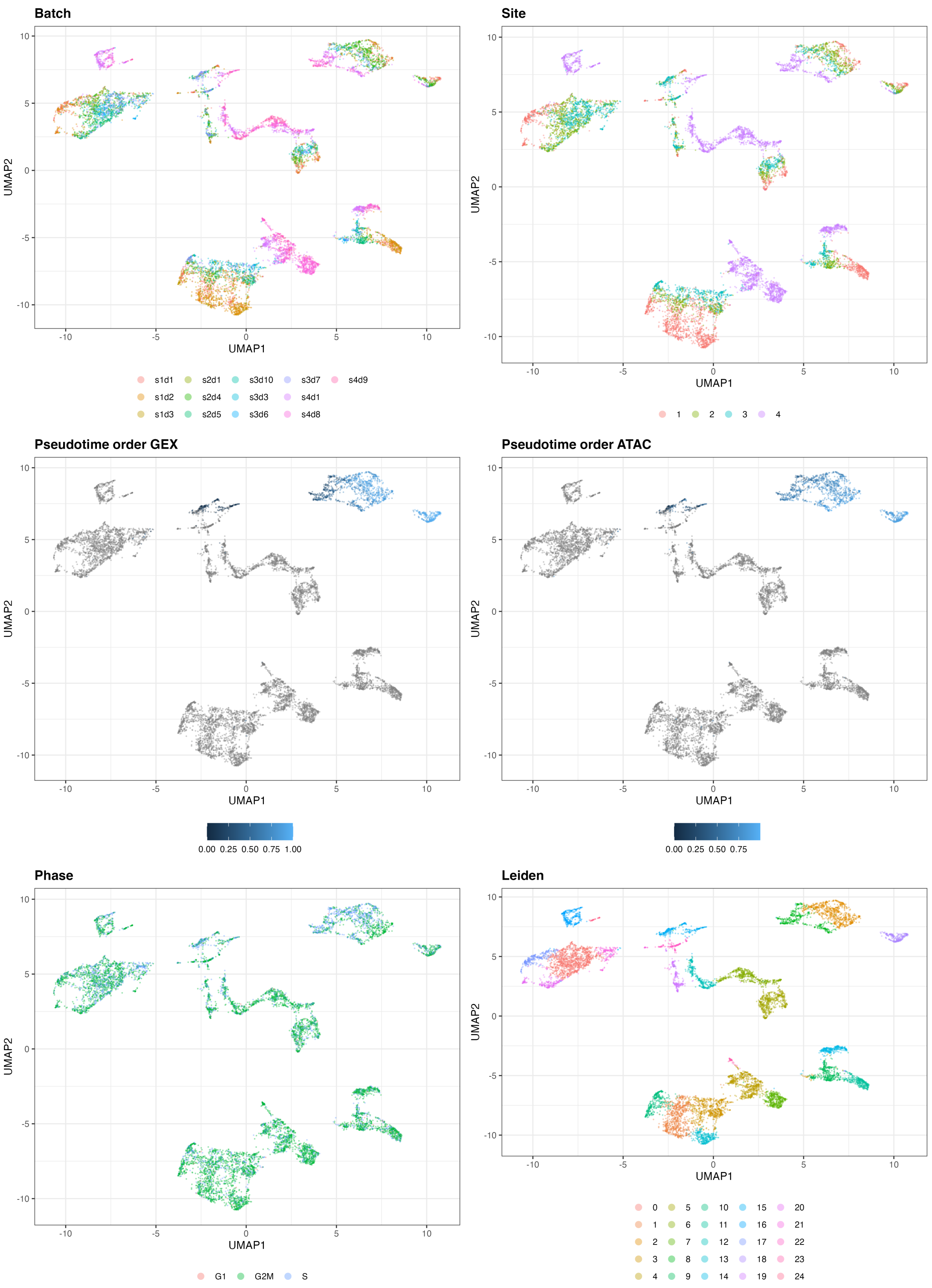


Figure 4: UMAP of the 10-dimensional latent space of Cobolt based on 10000 cells of one exemplary subsample from the Multiome dataset. The cells are color-coded by batch, site, pseudotime order GEX, pseudotime order ATAC, phase, and Leiden clustering.

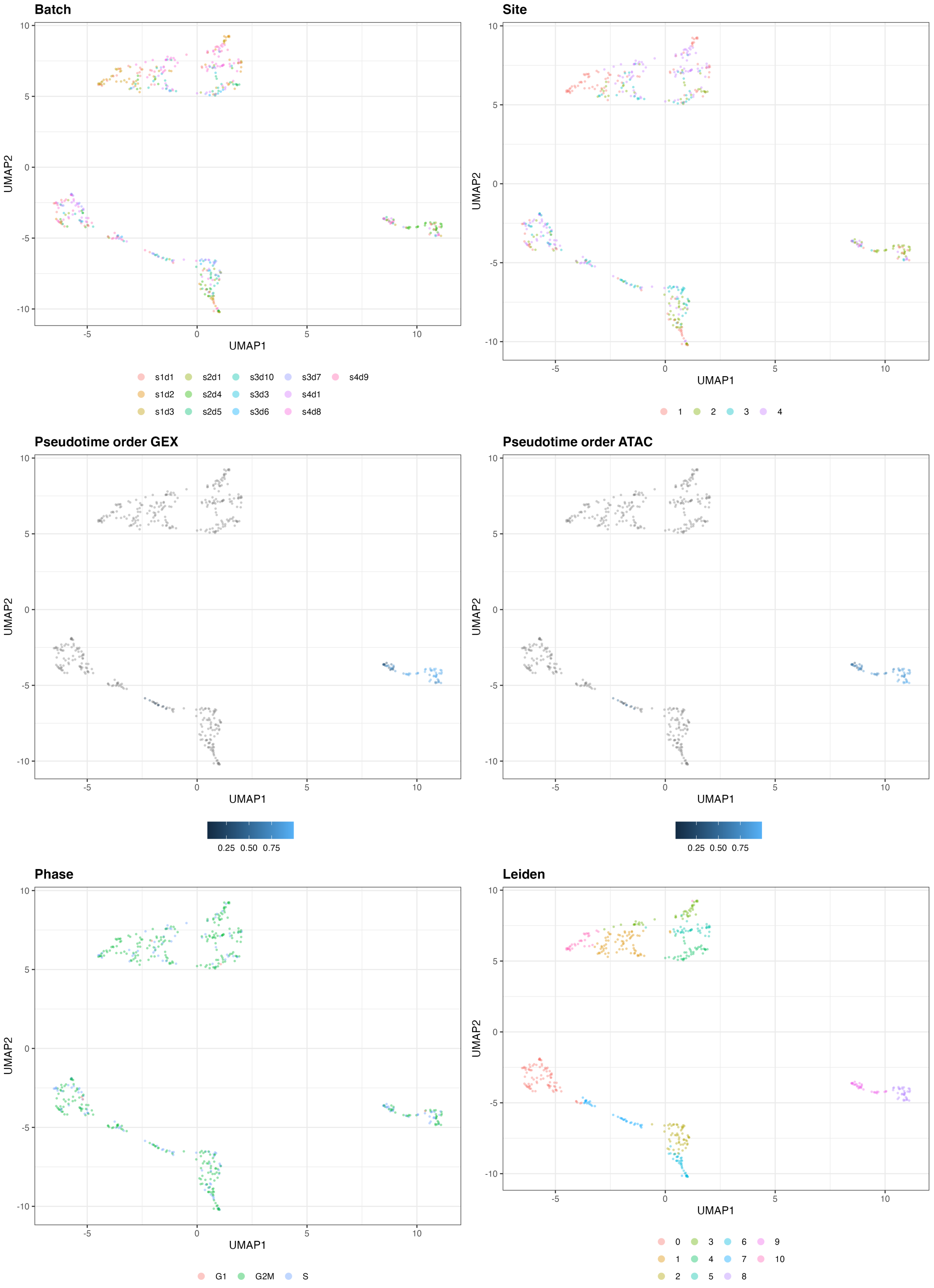


Figure 5: UMAP of the 10-dimensional latent space of scMM based on 500 cells of one exemplary subsample from the Multiome dataset. The cells are color-coded by batch, site, pseudotime order GEX, pseudotime order ATAC, phase, and Leiden clustering.

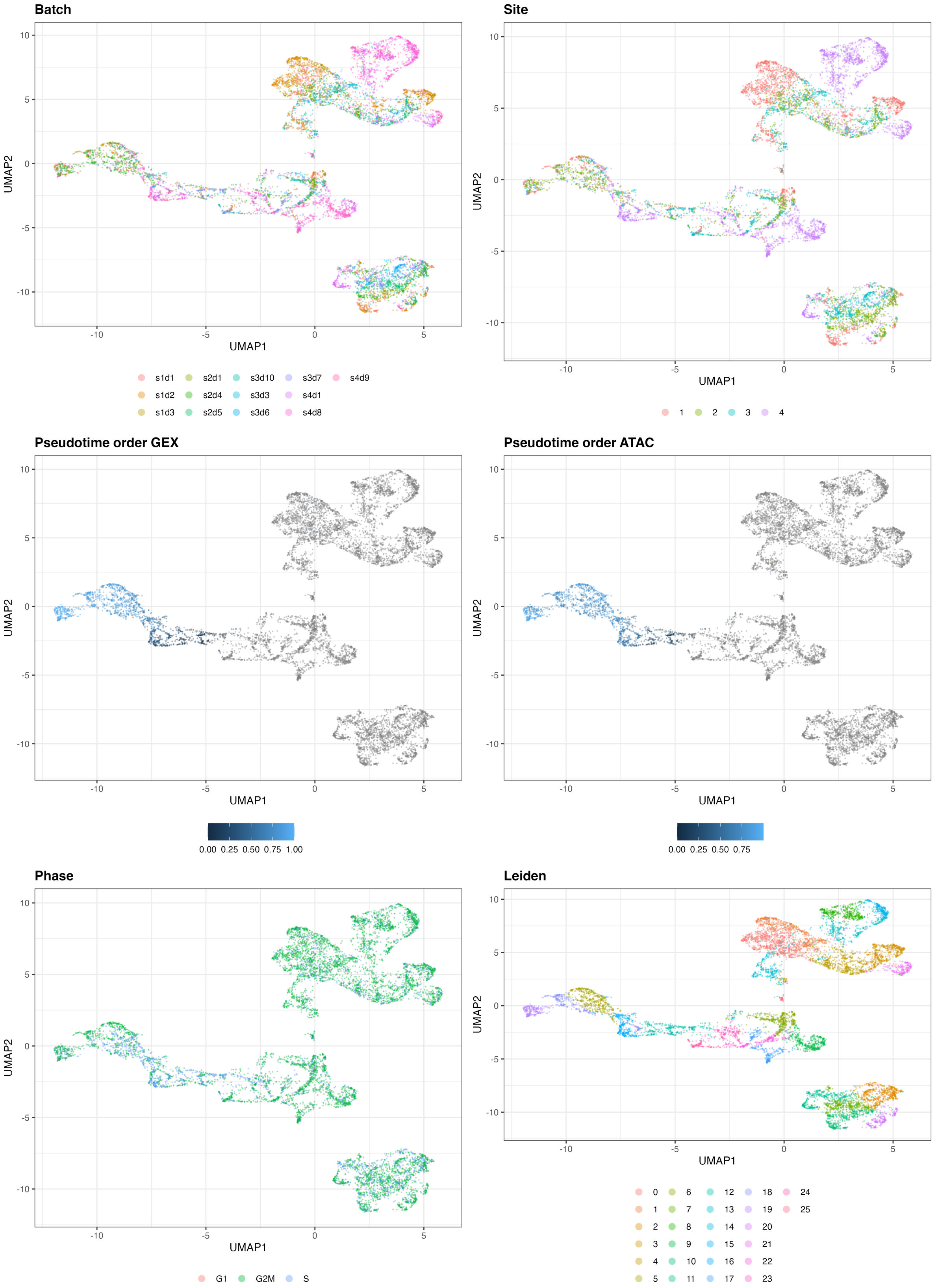


Figure 6: UMAP of the 10-dimensional latent space of scMM based on 10000 cells of one exemplary subsample from the Multiome dataset. The cells are color-coded by batch, site, pseudotime order GEX, pseudotime order ATAC, phase, and Leiden clustering.

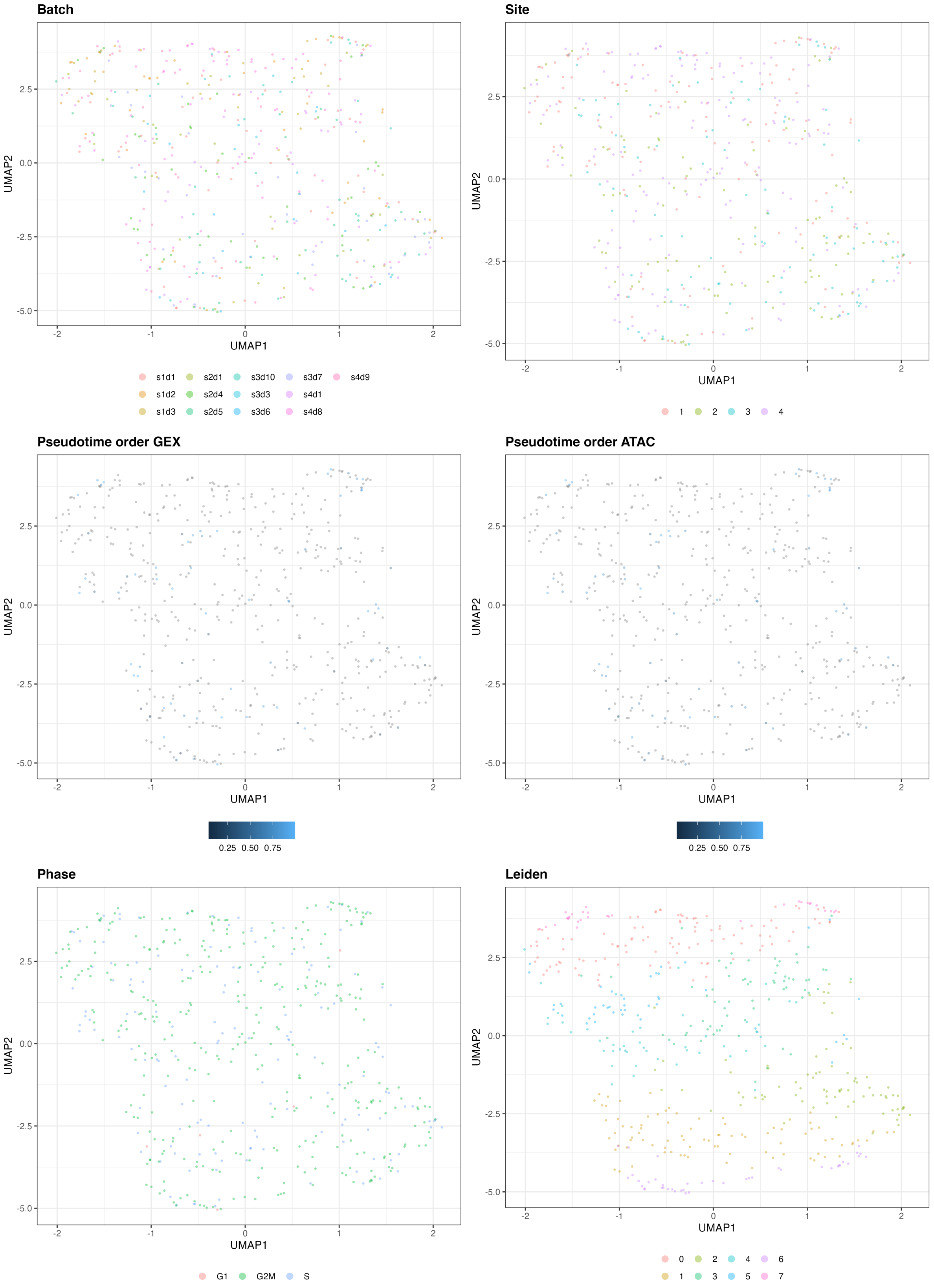


Figure 7: UMAP of the 10-dimensional latent space of MultiVI based on 500 cells of one exemplary subsample from the Multiome dataset. The cells are color-coded by batch, site, pseudotime order GEX, pseudotime order ATAC, phase, and Leiden clustering.

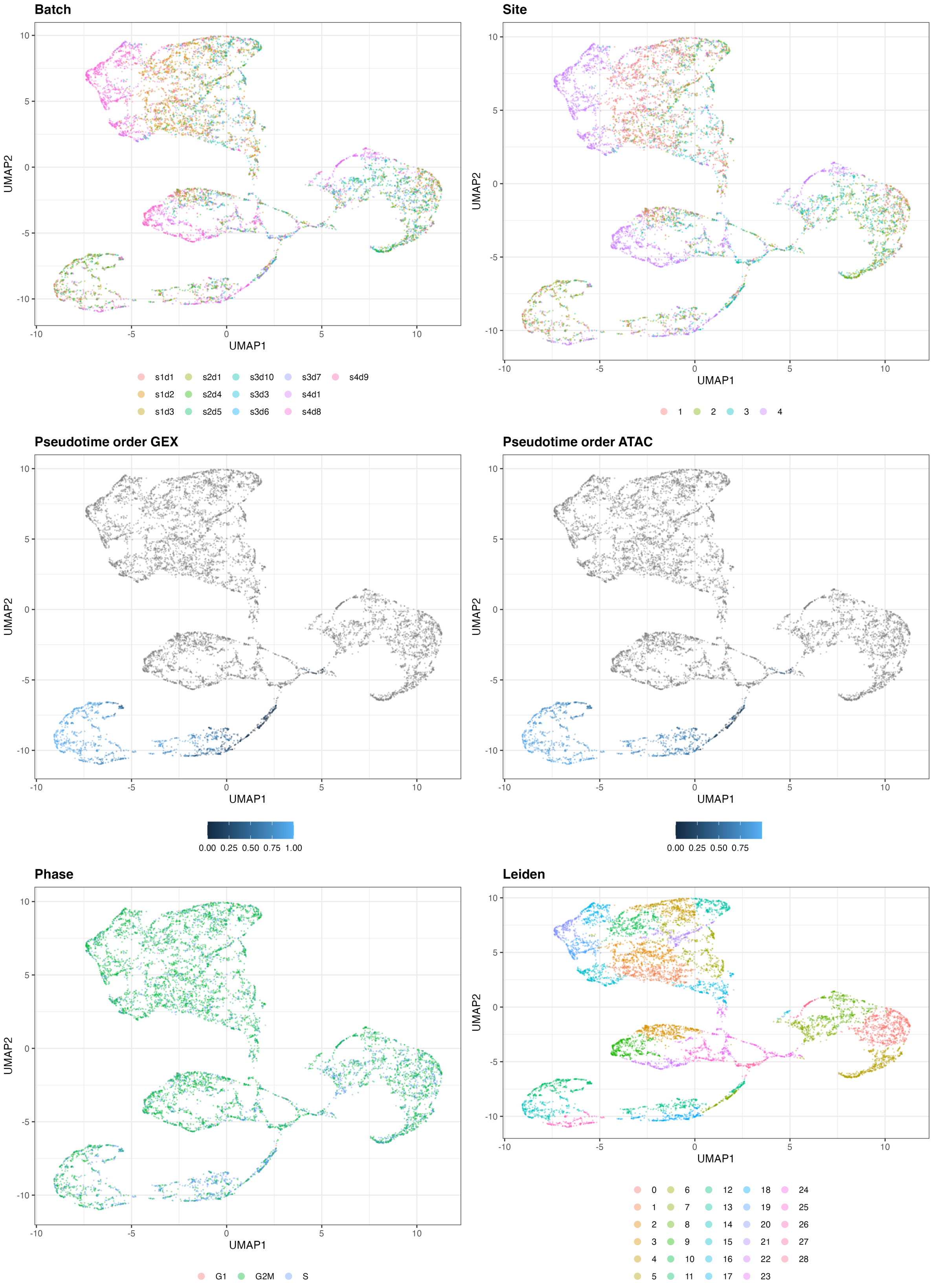


Figure 8: UMAP of the 10-dimensional latent space of MultiVI based on 10000 cells of one exemplary subsample from the Multiome dataset. The cells are color-coded by batch, site, pseudotime order GEX, pseudotime order ATAC, phase, and Leiden clustering.

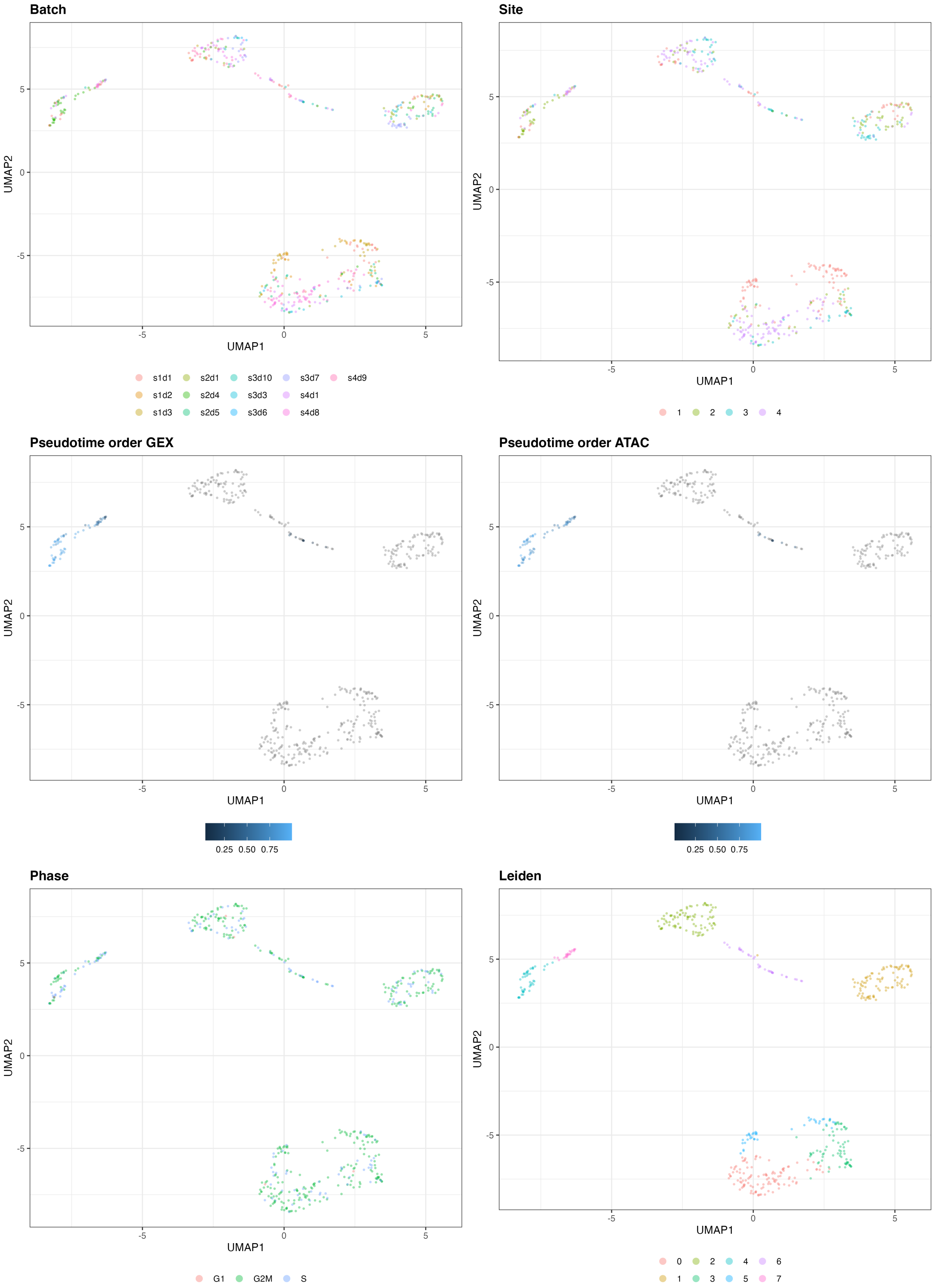


Figure 9: UMAP of the 10-dimensional latent space of scMVP based on 500 cells of one exemplary subsample from the Multiome dataset. The cells are color-coded by batch, site, pseudotime order GEX, pseudotime order ATAC, phase, and Leiden clustering.

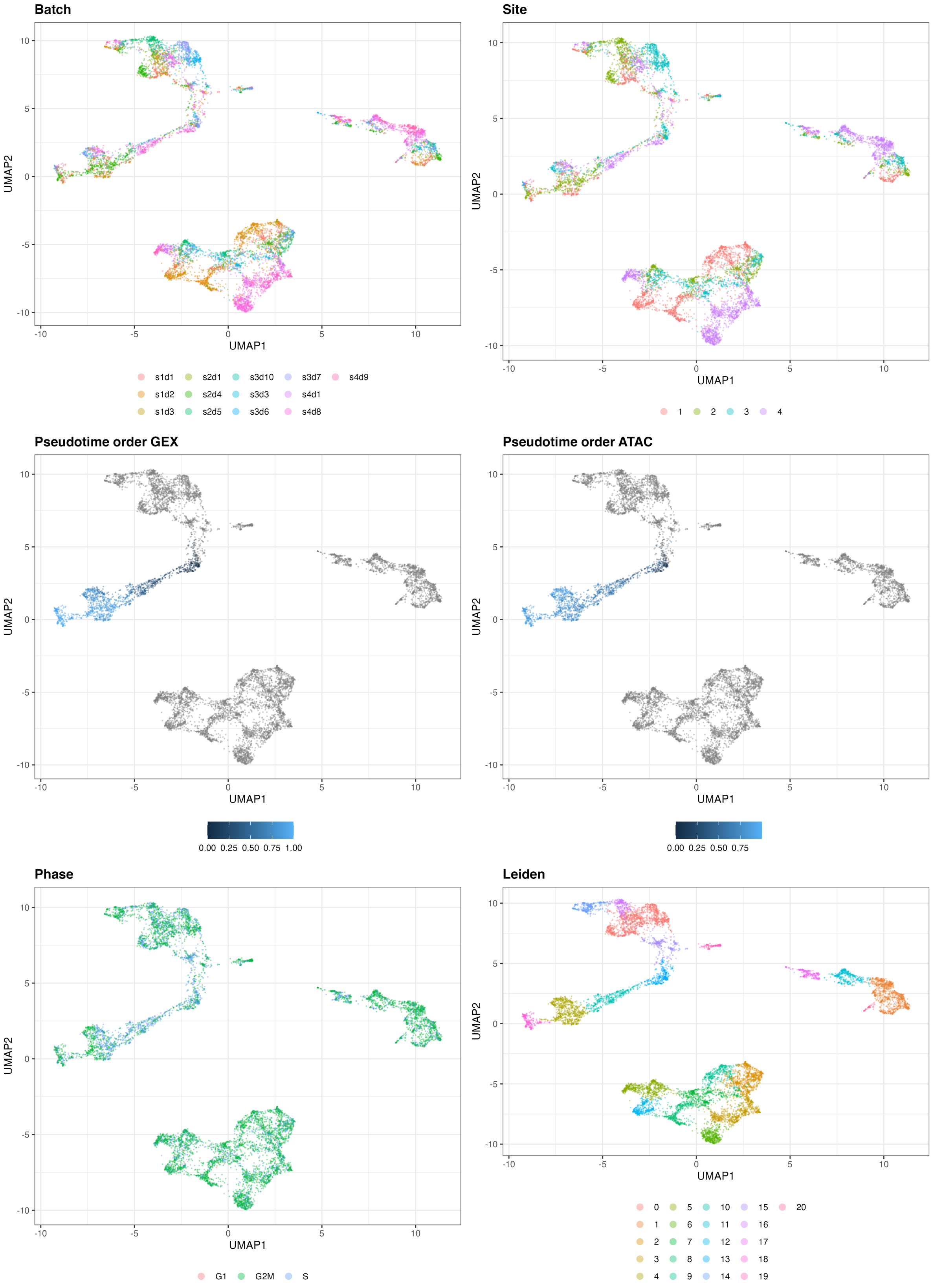


Figure 10: UMAP of the 10-dimensional latent space of scMVP based on 10000 cells of one exemplary subsample from the Multiome dataset. The cells are color-coded by batch, site, pseudotime order GEX, pseudotime order ATAC, phase, and Leiden clustering.

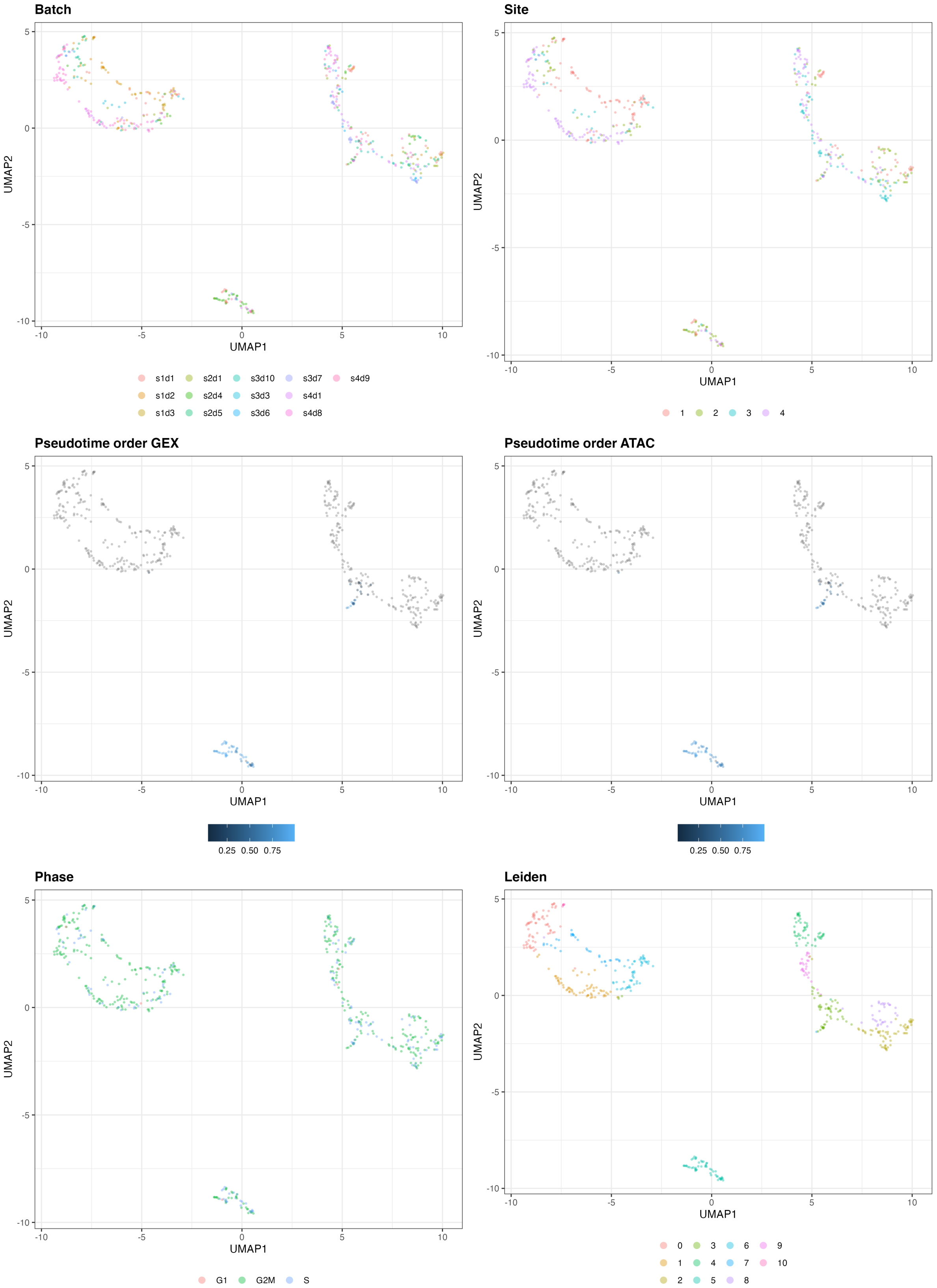


Figure 11: UMAP of the 10-dimensional latent space of DAVAE based on 500 cells of one exemplary subsample from the Multiome dataset. The cells are color-coded by batch, site, pseudotime order GEX, pseudotime order ATAC, phase, and Leiden clustering.

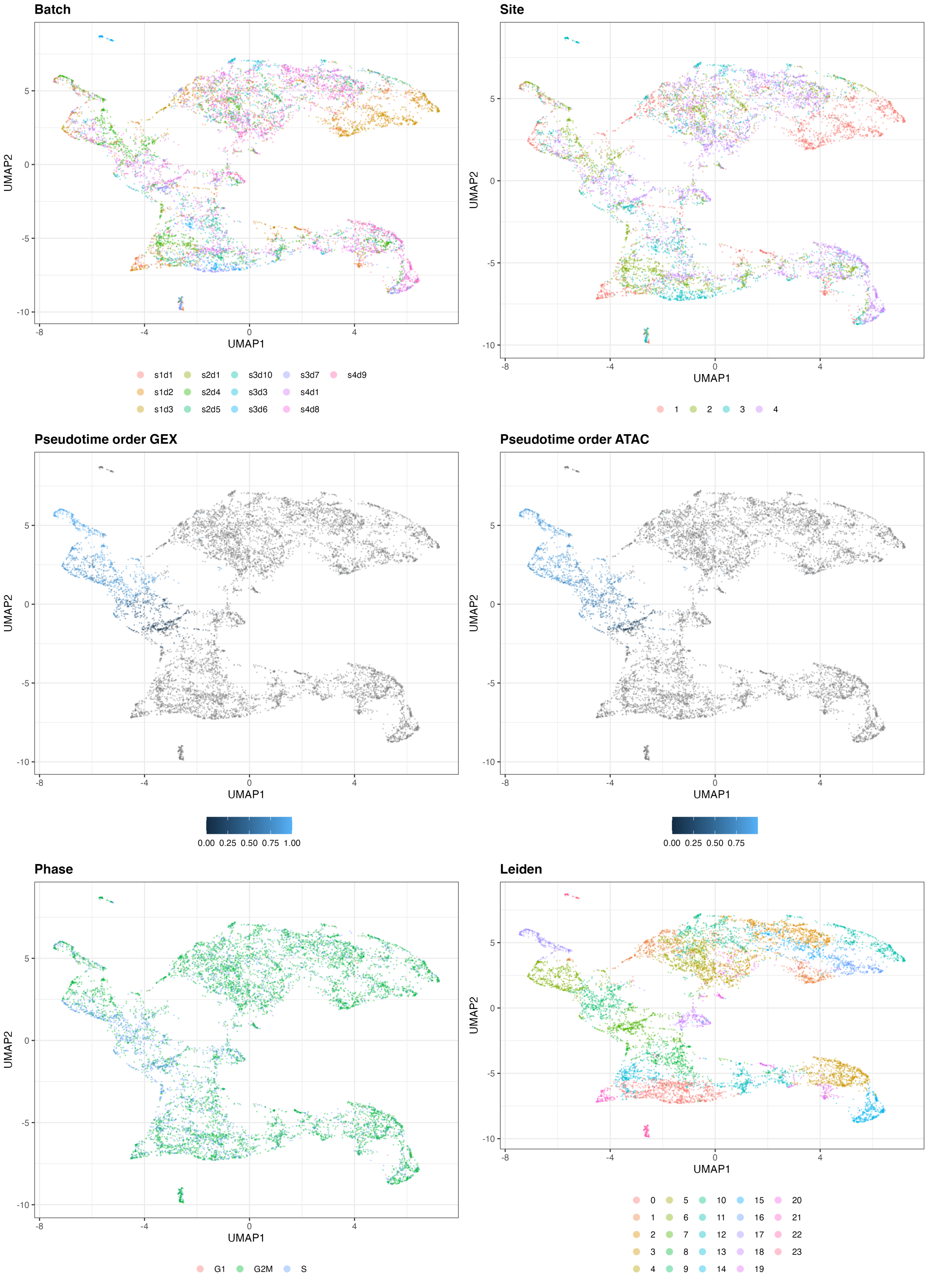


Figure 12: UMAP of the 10-dimensional latent space of DAVAE based on 10000 cells of one exemplary subsample from the Multiome dataset. The cells are color-coded by batch, site, pseudotime order GEX, pseudotime order ATAC, phase, and Leiden clustering.

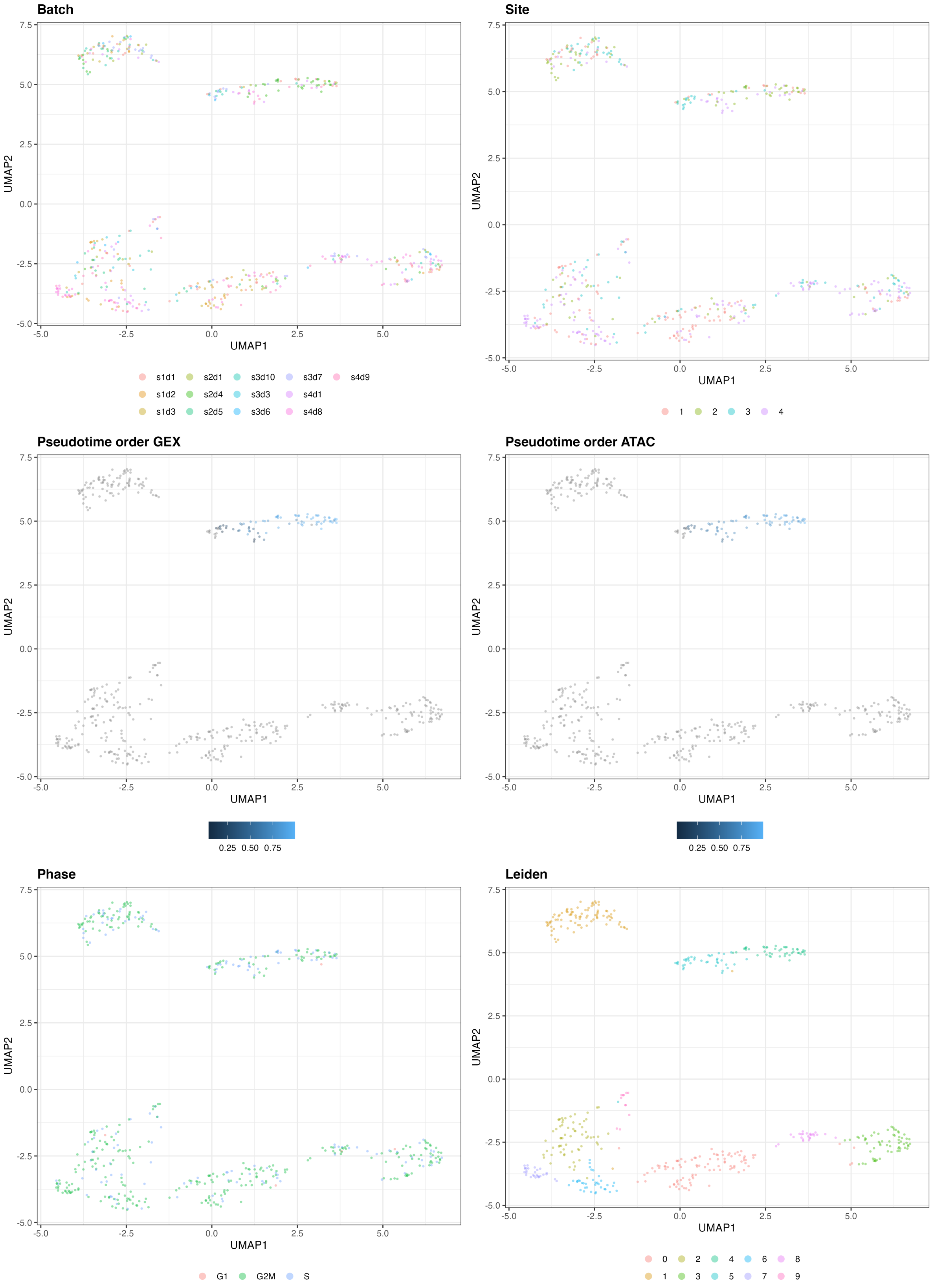


Figure 13: UMAP of the 10-dimensional latent space of Portal based on 500 cells of one exemplary subsample from the Multiome dataset. The cells are color-coded by batch, site, pseudotime order GEX, pseudotime order ATAC, phase, and Leiden clustering.

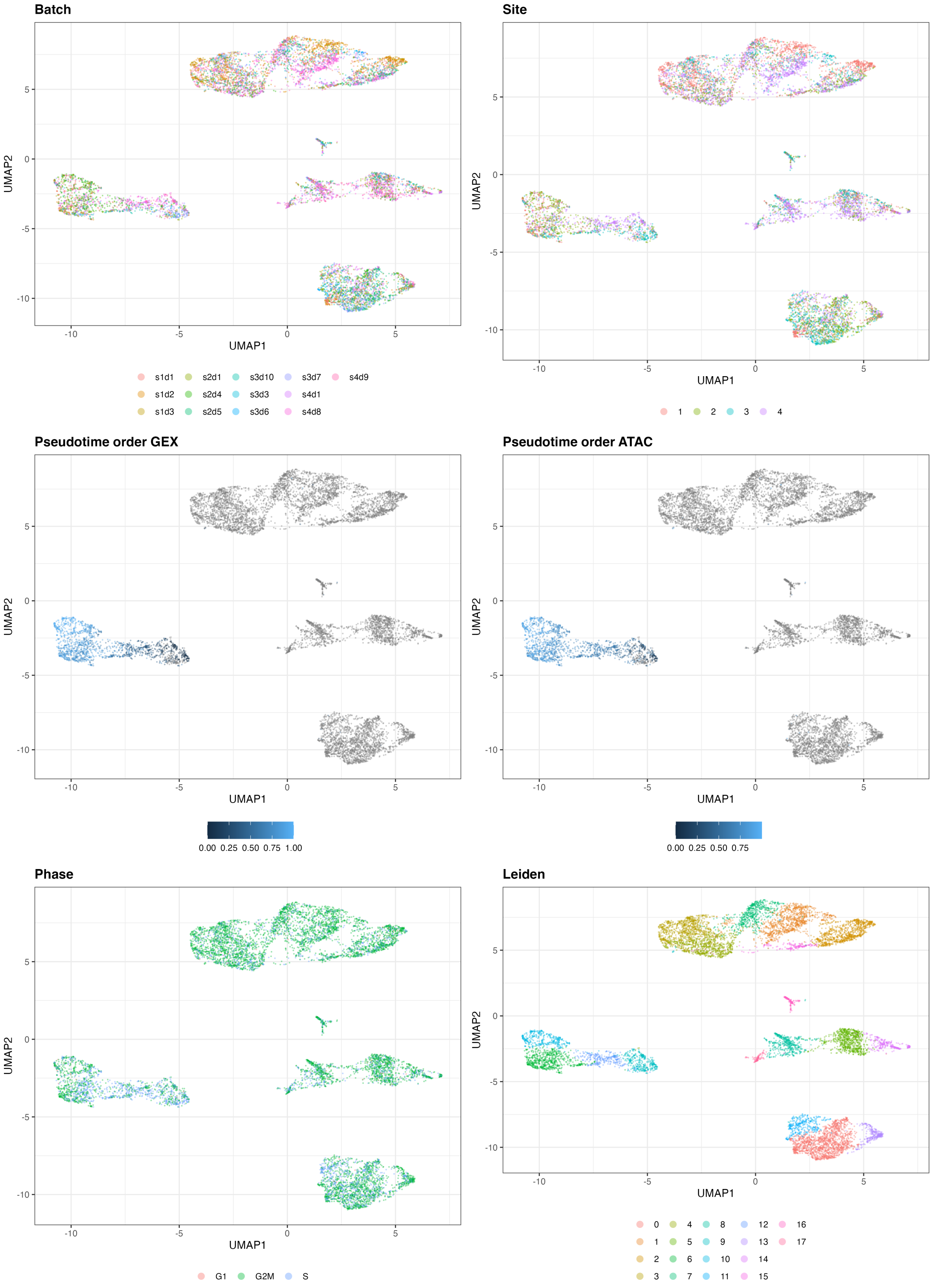


Figure 14: UMAP of the 10-dimensional latent space of Portal based on 10000 cells of one exemplary subsample from the Multiome dataset. The cells are color-coded by batch, site, pseudotime order GEX, pseudotime order ATAC, phase, and Leiden clustering.

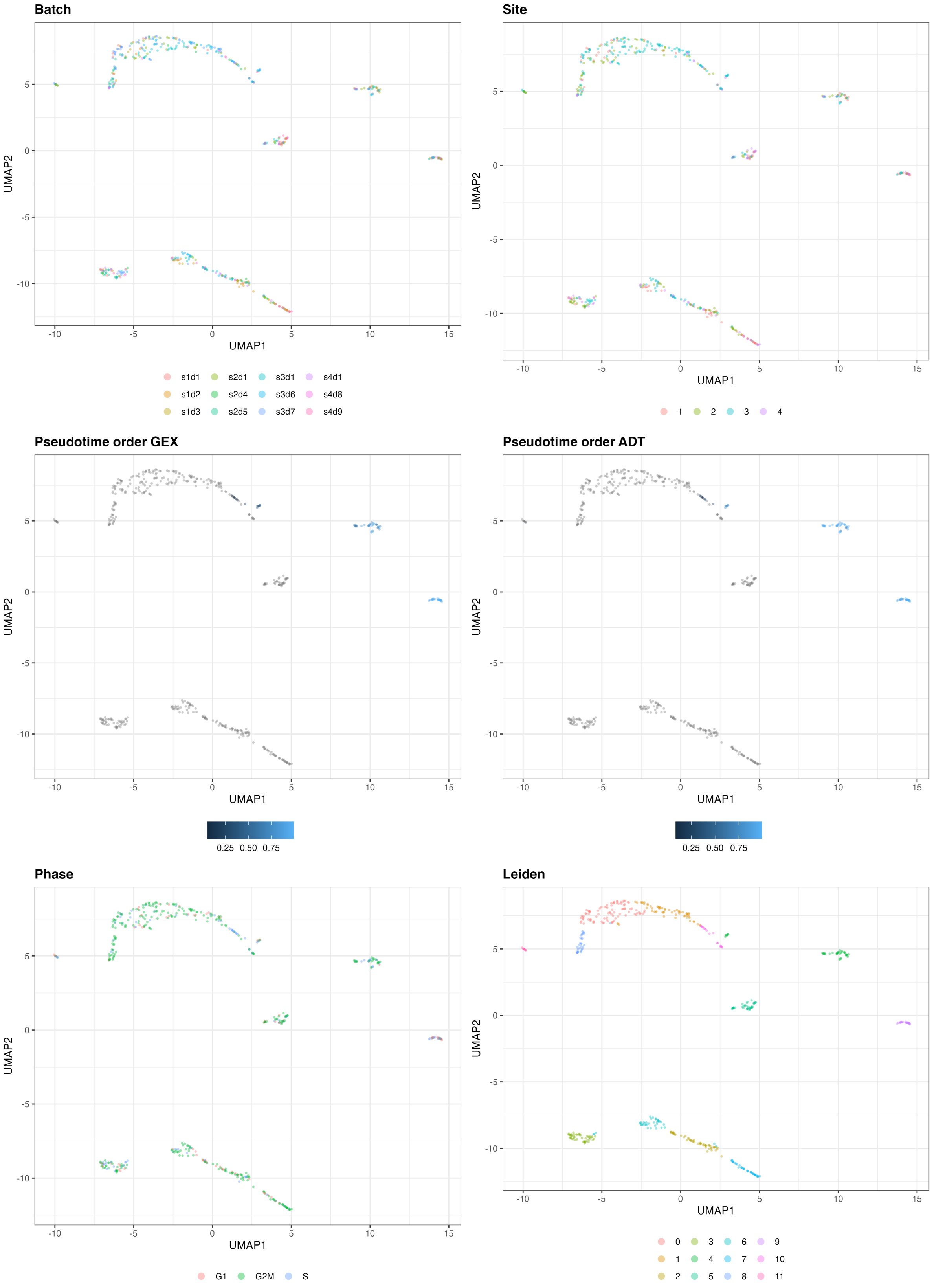


Figure 15: UMAP of the 10-dimensional latent space of Cobolt based on 500 cells of one exemplary subsample from the CITE-seq dataset. The cells are color-coded by batch, site, pseudotime order GEX, pseudotime order ADT, phase, and Leiden clustering.

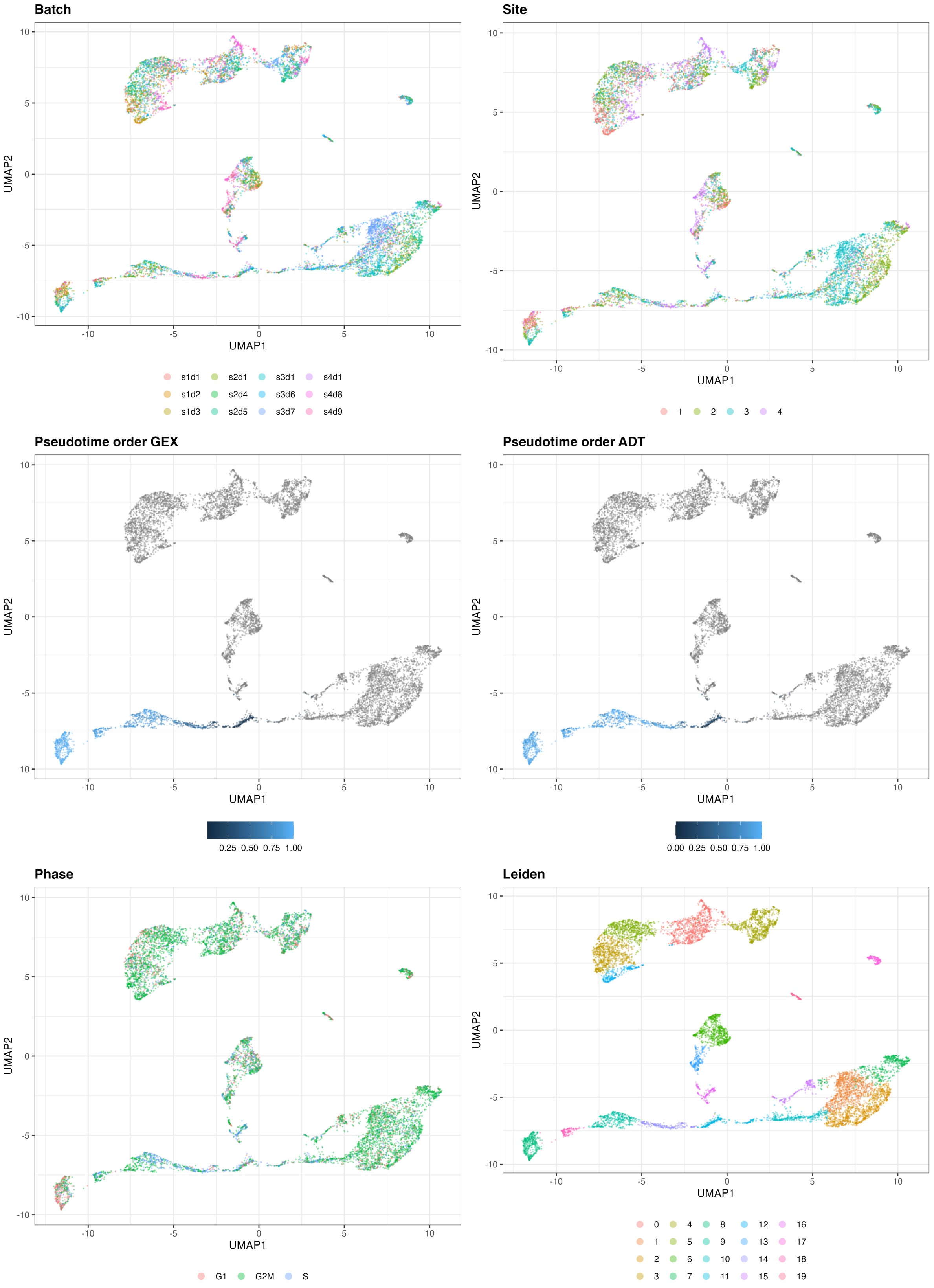


Figure 16: UMAP of the 10-dimensional latent space of Cobolt based on 10000 cells of one exemplary subsample from the CITE-seq dataset. The cells are color-coded by batch, site, pseudotime order GEX, pseudotime order ADT, phase, and Leiden clustering.

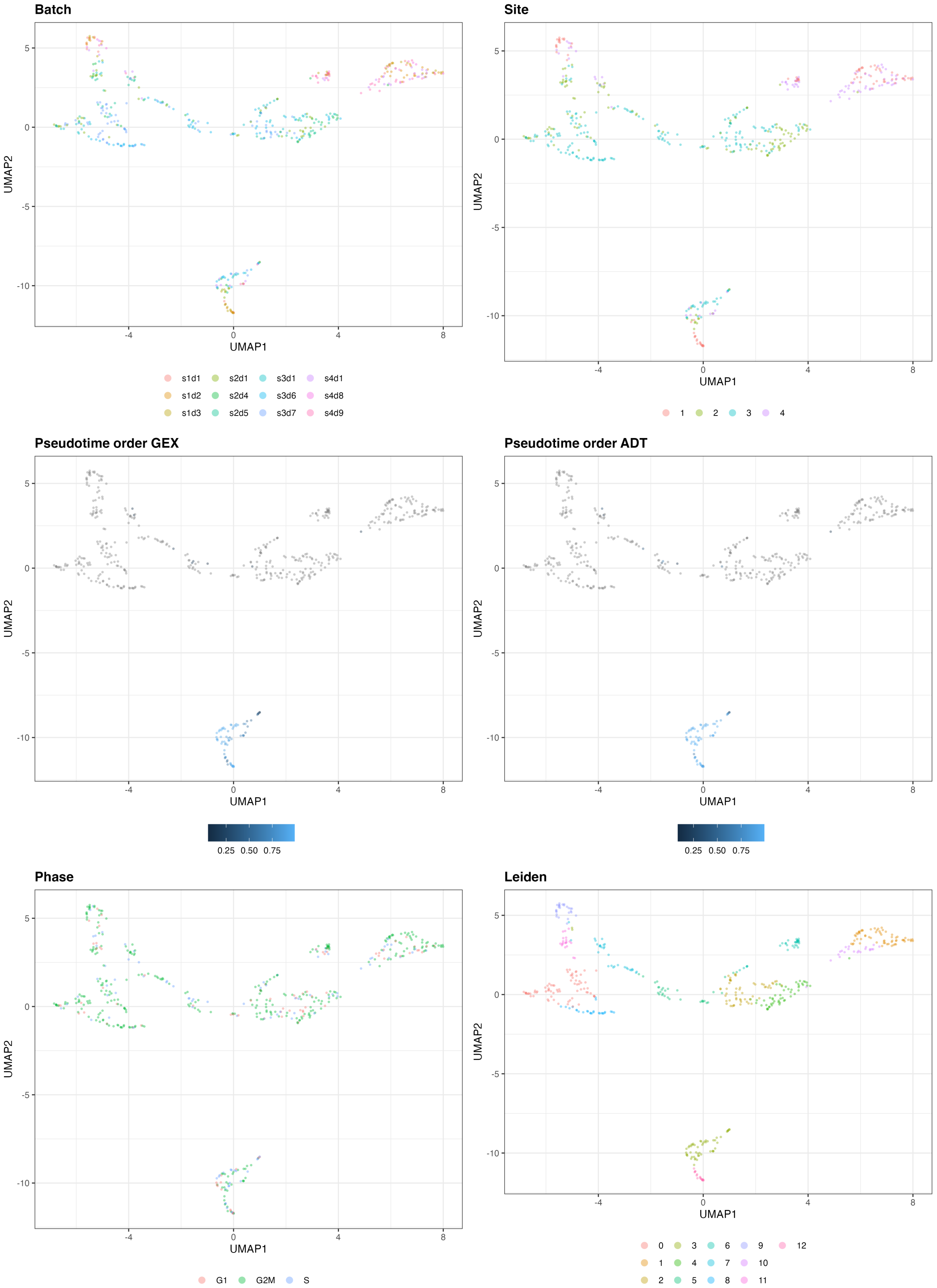


Figure 17: UMAP of the 10-dimensional latent space of scMM based on 500 cells of one exemplary subsample from the CITE-seq dataset. The cells are color-coded by batch, site, pseudotime order GEX, pseudotime order ADT, phase, and Leiden clustering.

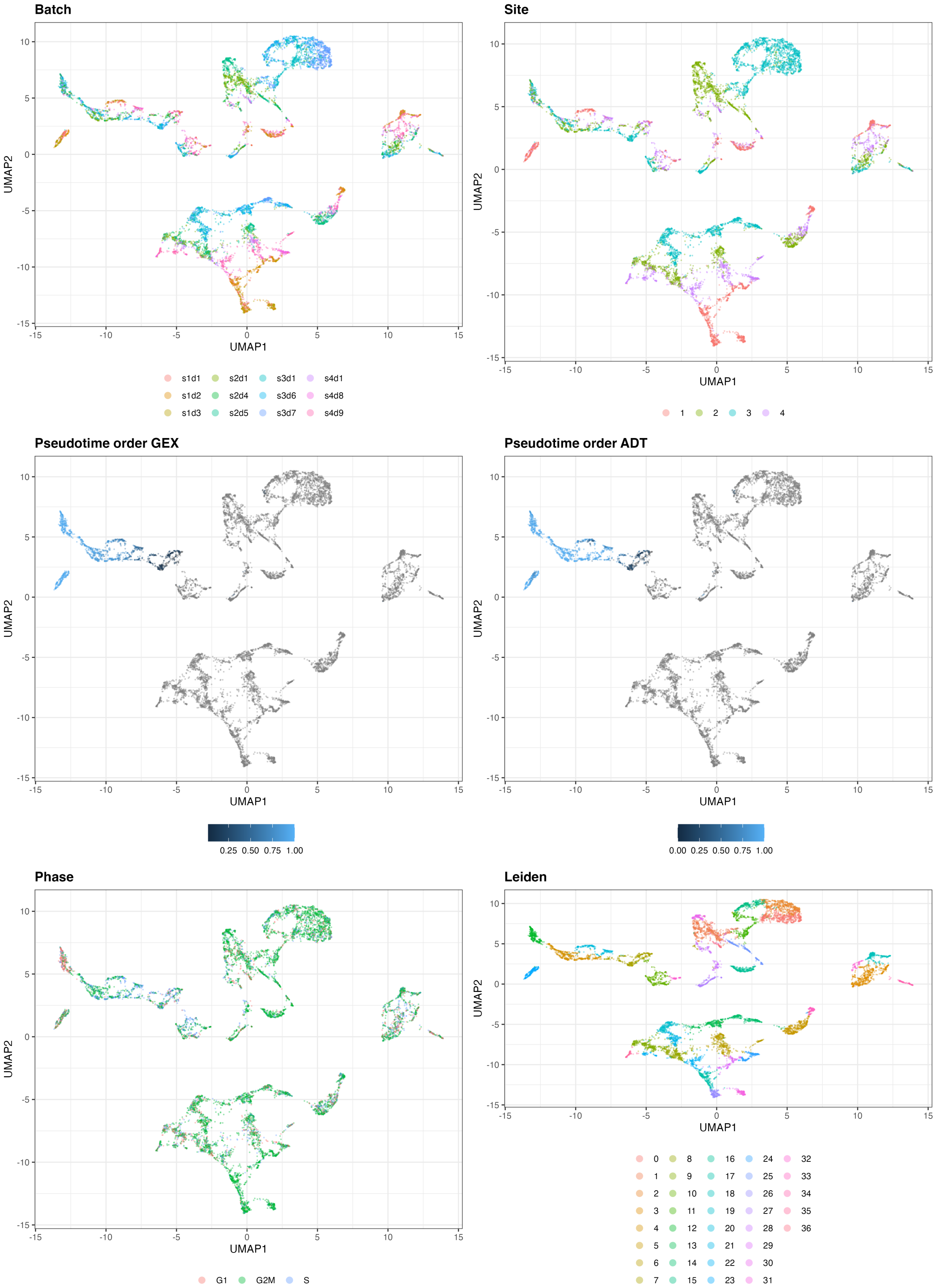


Figure 18: UMAP of the 10-dimensional latent space of scMM based on 10000 cells of one exemplary subsample from the CITE-seq dataset. The cells are color-coded by batch, site, pseudotime order GEX, pseudotime order ADT, phase, and Leiden clustering.

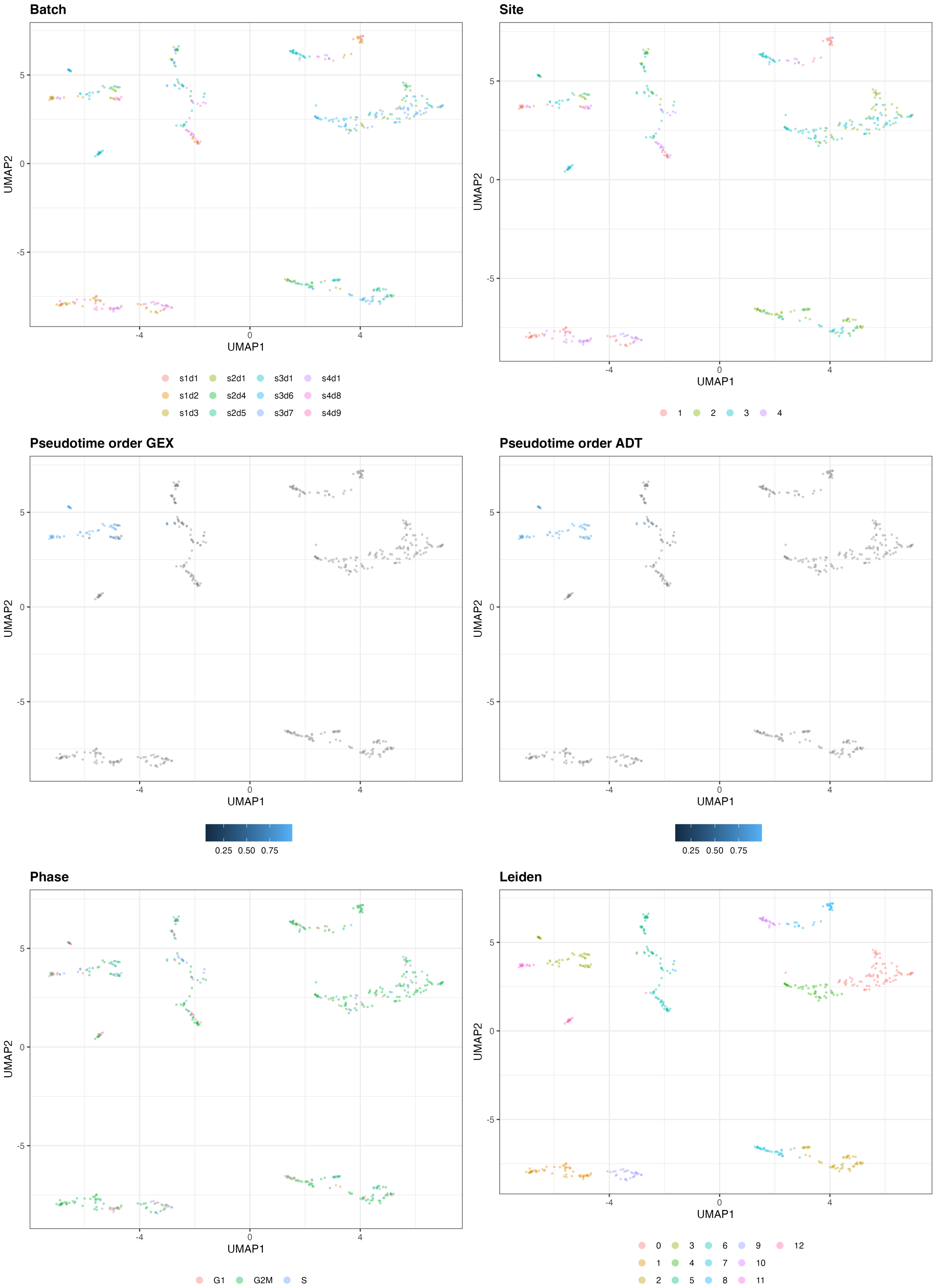


Figure 19: UMAP of the 10-dimensional latent space of TotalVI based on 500 cells of one exemplary subsample from the CITE-seq dataset. The cells are color-coded by batch, site, pseudotime order GEX, pseudotime order ADT, phase, and Leiden clustering.

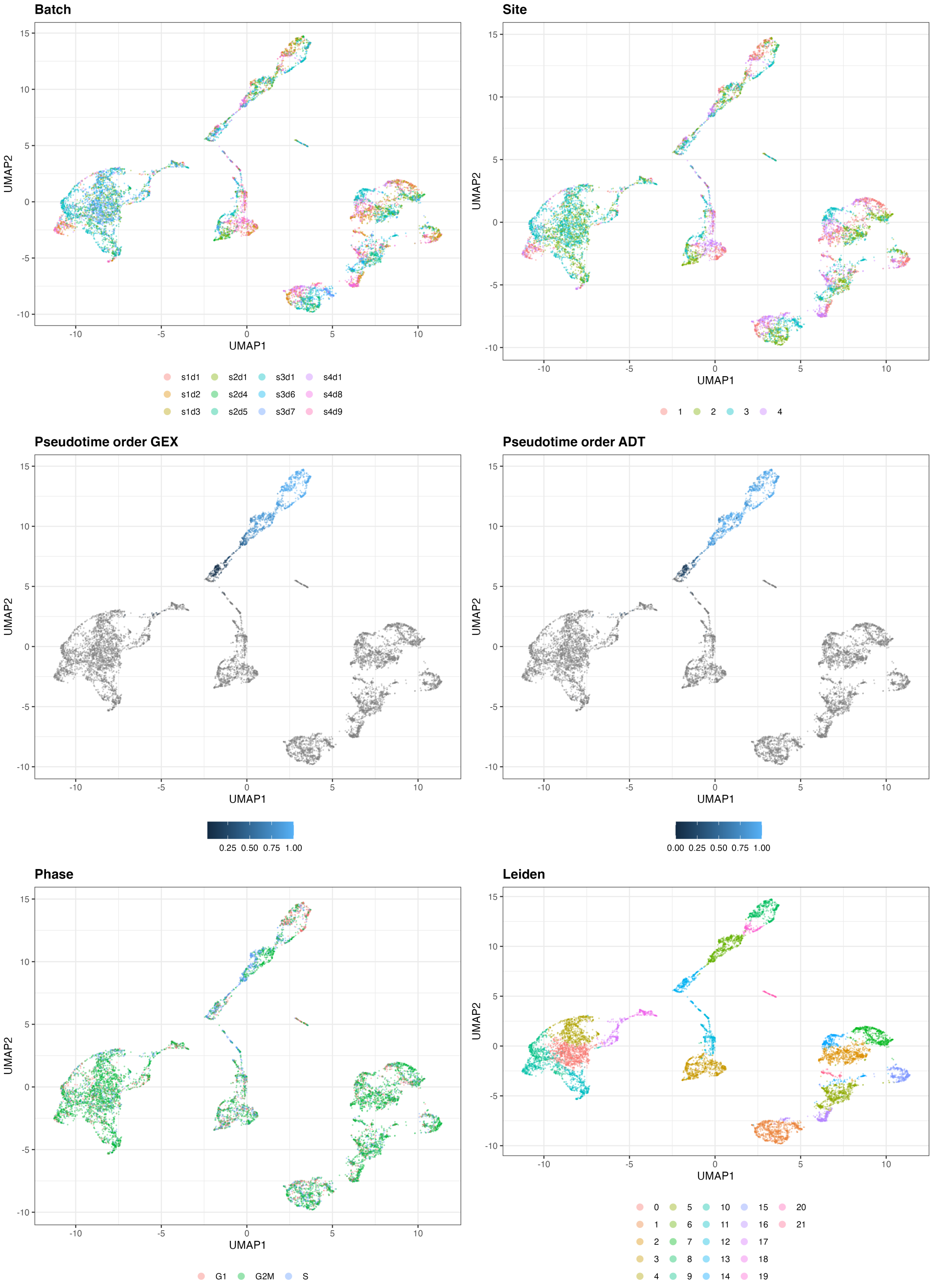


Figure 20: UMAP of the 10-dimensional latent space of TotalVI based on 10000 cells of one exemplary subsample from the CITE-seq dataset. The cells are color-coded by batch, site, pseudotime order GEX, pseudotime order ADT, phase, and Leiden clustering.

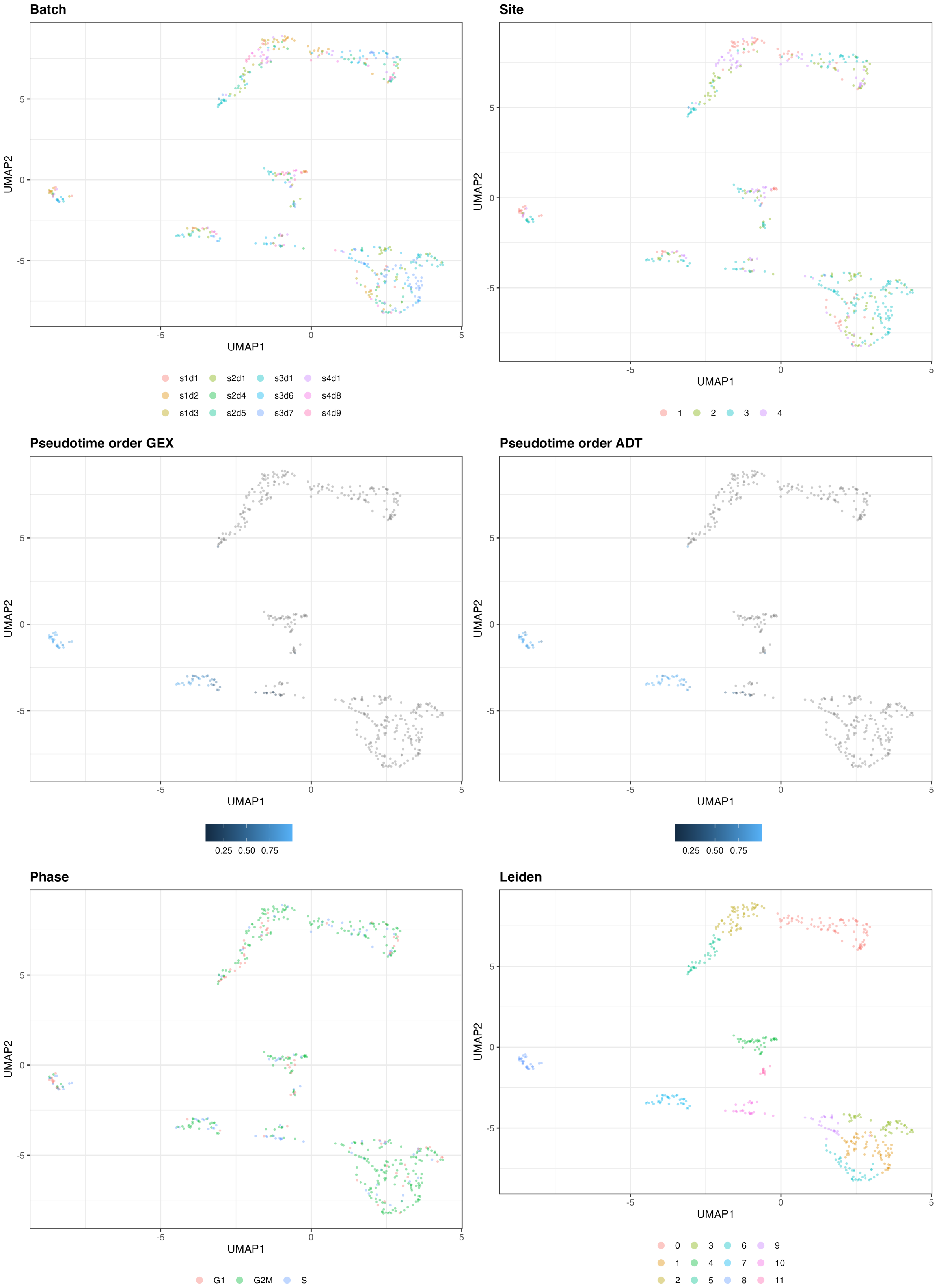


Figure 21: UMAP of the 10-dimensional latent space of SCALEX based on 500 cells of one exemplary subsample from the CITE-seq dataset. The cells are color-coded by batch, site, pseudotime order GEX, pseudotime order ADT, phase, and Leiden clustering.

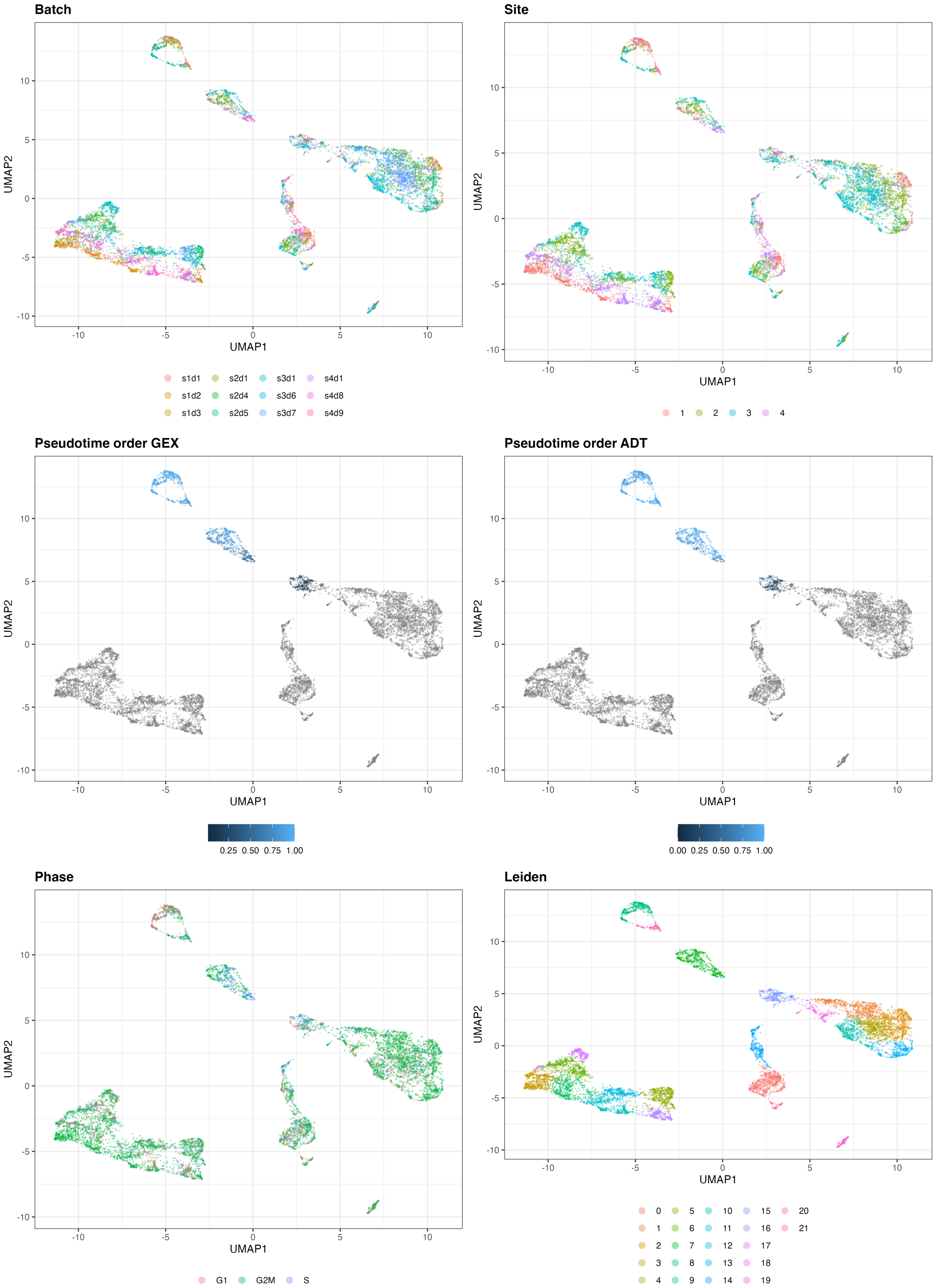


Figure 22: UMAP of the 10-dimensional latent space of SCALEX based on 10000 cells of one exemplary subsample from the CITE-seq dataset. The cells are color-coded by batch, site, pseudotime order GEX, pseudotime order ADT, phase, and Leiden clustering.

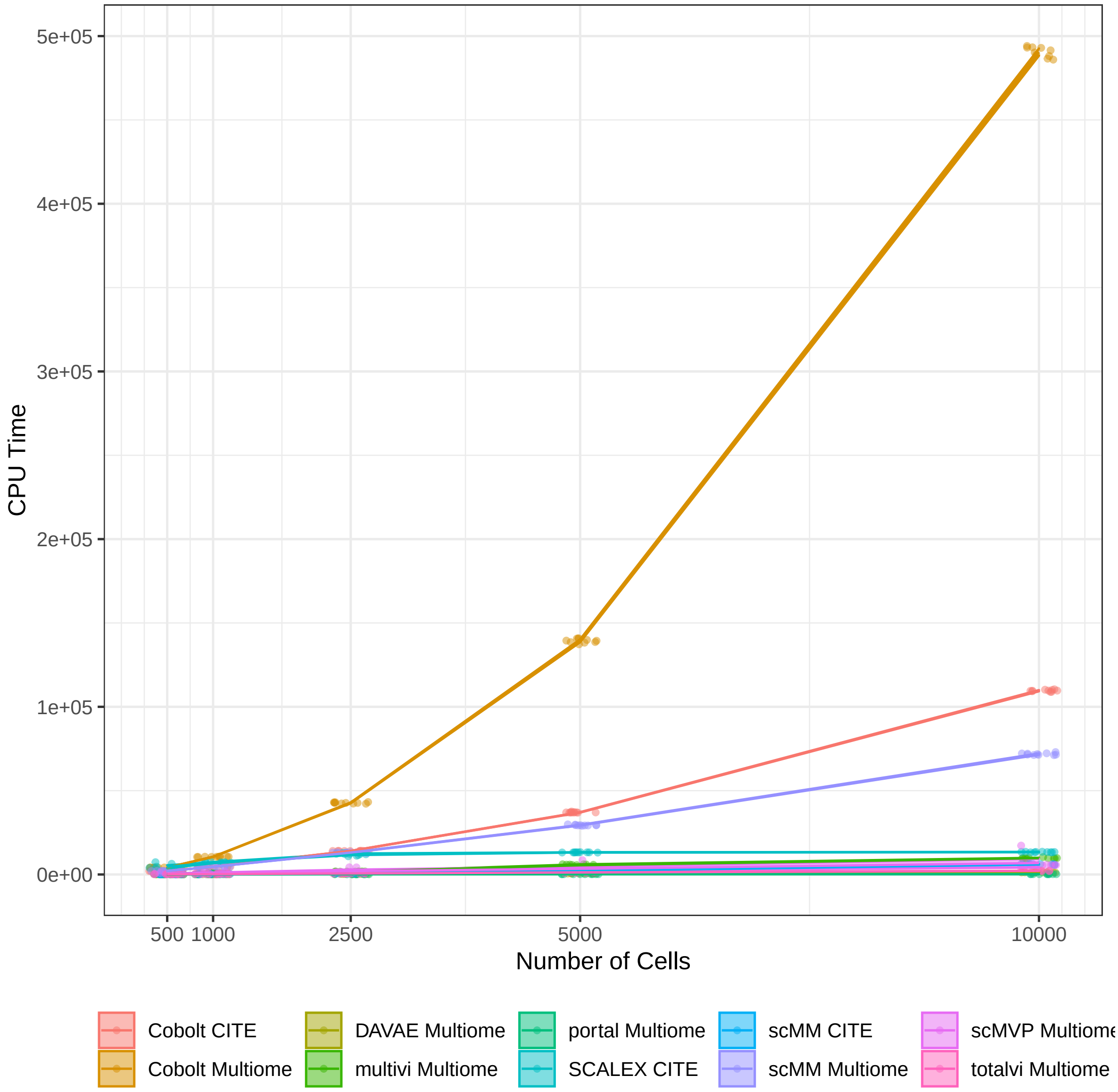


Figure 23: CPU time needed by each tool for processing the ten replicates of the five different cell numbers. For each tool, the increase in CPU is emphasized for each tool by a line, which runs through the mean of the CPU time of the ten replicates for each cell number condition. The ribbon indicates the 95% confidence interval.

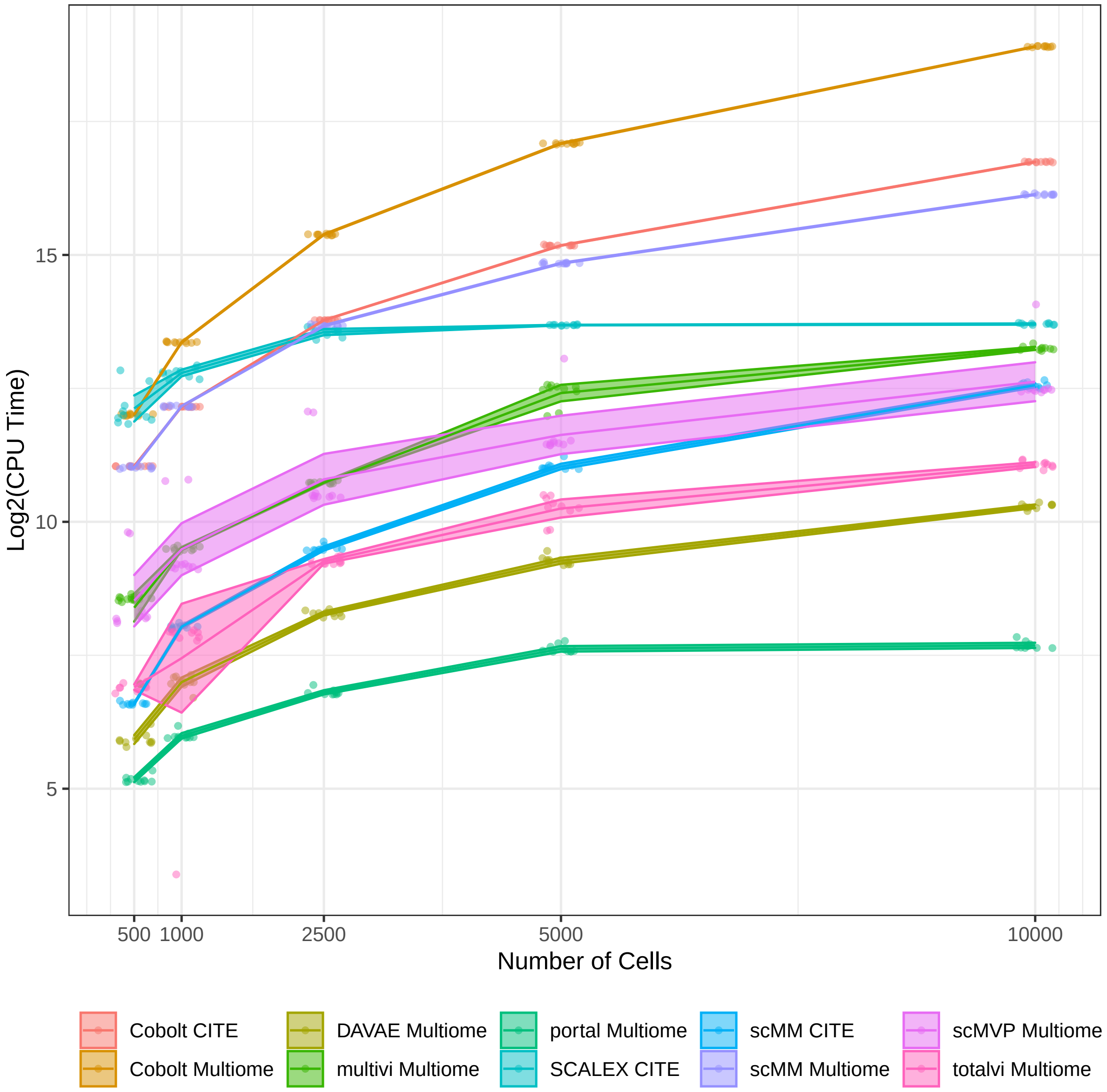


Figure 24: Log2(CPU time) needed by each tool for processing the ten replicates of the five different cell numbers. For each tool, the increase in CPU is emphasized for each tool by a line, which runs through the mean of the CPU time of the ten replicates for each cell number condition. The ribbon indicates the 95% confidence interval.

#### 2.2 Tables

Table 1: **Number of parameters per model.**

| Model | Numer of parameters |
|---|---|
| Cobolt (RNA + ATAC) | 4,314,758 |
| Cobolt (CITE-Seq) | 546,822 |
| scMM (RNA + ATAC) | 18,829,529 |
| scMM (CITE-Seq) | 2,827,908 |
| TotalVI | 4,760,917 |
| MultiVI | 19,849,578 |
| SCALEX | 4,295,040 |
| scMVP | 31,402,579 |
| DAVAE | 802,126 |
| Portal | 662,630 |

Table 2: **Spearman correlation between number of parameters and biological evaluation metrics.**

| **Biological Metrics** | | | | | |
|---|---|---|---|---|---|
| Metric | 500 Cells | 1000 Cells | 2500 Cells | 5000 Cells | 10,000 Cells |
| NMI | -0.405 | -0.503 | -0.454 | -0.356 | -0.209 |
| Cell Type ASW | -0.405 | -0.0614 | -0.0614 | -0.209 | 0.0614 |
| Trajectory Conservation | -0.681 | -0.595 | -0.405 | -0.0123 | -0.0614 |

Table 3: **Spearman correlation between number of parameters and technical evaluation metrics.**

| **Technical Metrics** | | | | | |
|---|---|---|---|---|---|
| Metric | 500 Cells | 1000 Cells | 2500 Cells | 5000 Cells | 10,000 Cells |
| Site ASW | -0.104 | -0.307 | -0.356 | -0.356 | -0.405 |
| Batch ASW | -0.215 | -0.399 | -0.399 | -0.399 | -0.399 |
| Graph Connectivity | -0.411 | -0.405 | -0.160 | -0.0123 | -0.0368 |
